# Extent of N-terminus exposure by altered long-range interactions of monomeric alpha-synuclein determines its aggregation propensity

**DOI:** 10.1101/740241

**Authors:** Amberley D. Stephens, Maria Zacharopoulou, Rani Moons, Giuliana Fusco, Neeleema Seetaloo, Anass Chiki, Philippa J. Hooper, Ioanna Mela, Hilal A. Lashuel, Jonathan J Phillips, Alfonso De Simone, Frank Sobott, Gabriele S. Kaminski Schierle

## Abstract

As an intrinsically disordered protein, monomeric alpha synuclein (aSyn) constantly reconfigures and probes the conformational space. Long-range interactions across the protein maintain its solubility and mediate this dynamic flexibility, but also provide residual structure. Certain conformations lead to aggregation prone and non-aggregation prone intermediates, but identifying these within the dynamic ensemble of monomeric conformations is difficult. Herein, we used the biologically relevant calcium ion to investigate the conformation of monomeric aSyn in relation to its aggregation propensity. By using calcium to perturb the conformational ensemble, we observe differences in structure and intra-molecular dynamics between two aSyn C-terminal variants, D121A and pS129, and the aSyn familial disease mutants, A30P, E46K, H50Q, G51D, A53T and A53E, compared to wild-type (WT) aSyn. We observe that the more exposed the N-terminus and the beginning of the NAC region are, the more aggregation prone monomeric aSyn conformations become. N-terminus exposure occurs upon release of C-terminus interactions when calcium binds, but the level of exposure is specific to the aSyn mutation present. There was no correlation between single charge alterations, calcium affinity, or the number of ions bound on aSyn’s aggregation propensity, indicating that sequence or post-translation modification (PTM)-specific conformational differences between the N- and C-termini and the specific local environment mediate aggregation propensity instead. Understanding aggregation prone conformations of monomeric aSyn and the environmental conditions they form under will allow us to design new therapeutics targeted to the monomeric protein, to stabilise aSyn in non-aggregation prone conformations, by either preserving long-range interactions between the N- and C-termini or by protecting the N-terminus from exposure.

## Introduction

In Parkinson’s disease (PD) and other synucleinopathies, the monomeric protein alpha synuclein (aSyn) becomes destabilised, misfolds and aggregates into insoluble, highly structured and β-sheet containing fibrils which form part of Lewy bodies (LB) and Lewy neurites (LN) ^1,2^. In its monomeric form, aSyn is a soluble, intrinsically disordered protein (IDP) that is highly flexible and thereby enables plasticity in its function. In particular, it has been proposed that aSyn plays a role in synaptic vesicle recycling and homeostasis in neurons ^3,4^. Transient and dynamic electrostatic and hydrophobic intramolecular interactions maintain aSyn in its soluble monomeric form. These intramolecular interactions are responsible for aSyn’s smaller radius of gyration (*Rg*) than expected for a 140 amino acid residue (aar) unfolded protein and suggest that some residual structure remains ^5^. Therefore, the word ‘monomer’ actually describes a plethora of conformational states which are constantly reconfiguring. These dynamic interactions are heavily influenced by the surrounding environment of aSyn and their disruption can lead to skewing of the ensemble of monomeric conformations ^6^. This, in turn, may influence which aggregation competent/incompetent pathways are taken and whether these are subsequently toxic or not. Identifying conformations of aggregation prone monomeric aSyn or the environment that can destabilise monomeric conformations to favour aggregation will aid in the design of anti-aggregation therapeutics to stabilise the native aSyn conformation.

aSyn is a characteristic IDP with high opposing charge at its termini and low overall hydrophobicity ^5^. Monomeric aSyn has three characteristic main regions; the N-terminus, aar 1-60, which is overall positively charged, the non-amyloid-β component (NAC) region, aar 61-89, is hydrophobic and forms the core of fibrils during aggregation ^7^, and the C-terminus, aar 90-140, is a highly negatively charged region which binds metal ions ^8^ (Figure 1a). To date, six disease-related mutations have been identified in the *SNCA* gene, encoding the aSyn protein, A30P, E46K, H50Q, G51D, A53T, A53E, which are a hallmark for hereditary autosomal dominant PD and are primarily linked to early age but also late age of onset (H50Q) ^9–15^. However, genetic mutations and multiplications of the *SNCA* gene only account for 5-10% of PD cases and the remaining cases are sporadic (idiopathic) and age-related ^16^. Yet, we still have not identified mechanistically why these mutations lead to early-onset PD, or what triggers misfolding of WT aSyn.

**Figure 1.**
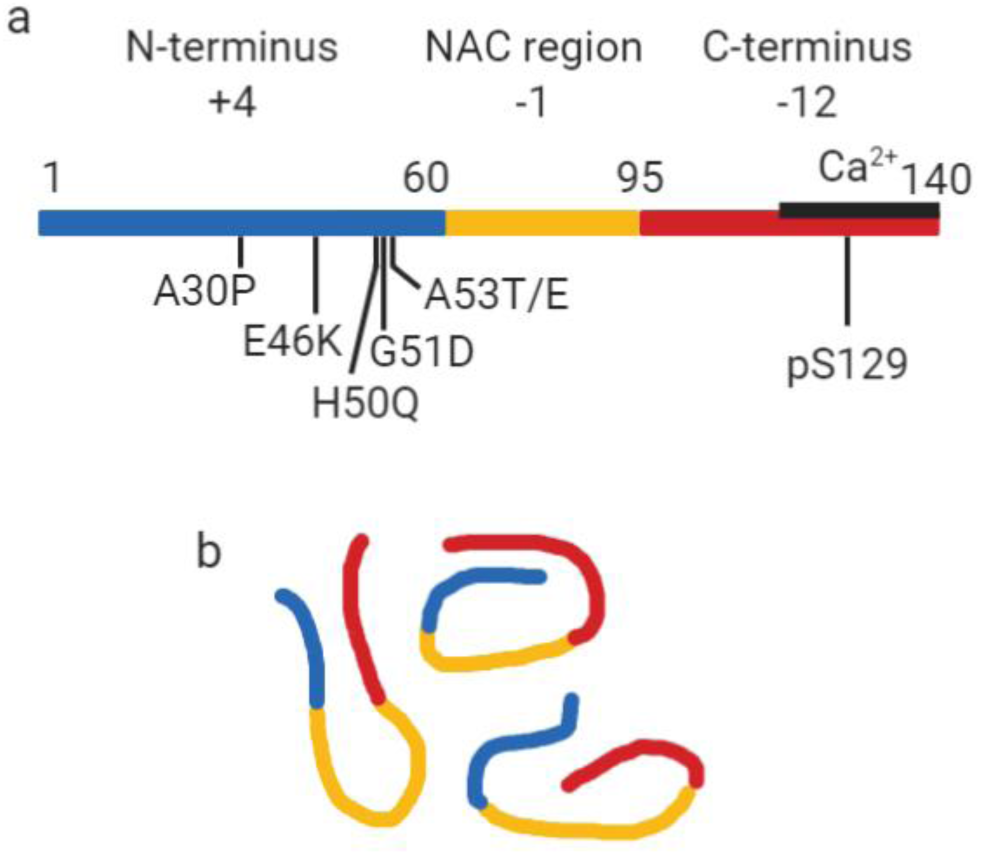
Representation of the regions of monomeric aSyn. (a) Monomeric aSyn is defined by three regions, the N-terminus, aar 1-60 (blue) with an overall charge of +4, contains the familial mutations A30P, E46K, H50Q, G51D, A53E and A53T. The non-Amyloid-β component (NAC) region, aar 61-95 (yellow), has an overall charge of -1, is highly hydrophobic and forms the core of fibrils. The C-terminus, aar 96-140 (red), is highly negatively charged with an overall charge of -12. Residue S129 is commonly phosphorylated (pS129) in Lewy bodies, but rarely in soluble aSyn. The calcium binding region (black line) is also found at the C-terminus and spans residues 115-140. (b) Monomeric aSyn is highly dynamic and visits a large conformational space. Interactions between the N- and C-termini and NAC region maintain it in a soluble form.

Intramolecular long-range interactions of aSyn have been detected between many different regions of aSyn. Electrostatic interactions, mediated by the positively charged N-terminus and negatively charged C-terminus, as well as hydrophobic interactions between some residues of the C-terminus and NAC region of aSyn, have been identified using a range of techniques including different nuclear magnetic resonance (NMR) techniques, mass spectrometry (MS) and hydrogen-deuterium exchange MS (HDX-MS) ^17–25^ (for a review see ^6^) (Figure 1b). The importance of these long-range interactions was demonstrated in studies in which charge and hydrophobicity of the protein were altered by mutations, particularly at the C-terminus, leading to differences in aSyn’s aggregation propensity ^2,26–28^. Reduction of charge also occurs during the binding of metal ions, salt ions or polyamines which leads to shielding of the charged N- and C-termini and which permits more energetically favourable packing ^8,29^.

Furthermore, post-translational modifications (PTM), such as nitration and phosphorylation, also alter aggregation rates of aSyn. In particular, phosphorylation of S129 which increases the negative charge of the C-terminus by the addition of a PO^4-^ group seems to be pertinent in disease as only 4% of monomeric aSyn is phosphorylated, yet 96% of aSyn in LB and LN are phosphorylated ^30^. However, it is not clear whether phosphorylation of S129 is involved in the physiological or pathological function of aSyn, whether it enhances aggregation ^31,32^ or retards aggregation ^33^. In terms of disease association, the presence of aSyn familial mutations leads to different aggregation rates dependent on the mutation. NMR experiments have shown that C-terminus residues are transiently in contact with all six mutation sites at the N-terminus via long-range interactions ^23^, yet the different mutations lead to differences in levels of solvent exposure, destabilisation, perturbation of the ensemble of conformers and alterations in long-range interactions ^34–37^. Identifying conformations or significant long-range interactions that maintain soluble aSyn is critical in understanding what triggers aSyn misfolding, but the difficulty in identifying these long-range interactions and determining the influence of mutations and aggregation prone conformations of aSyn lies in the complexity of sampling an ensemble of dynamic conformations of the monomeric protein.

In the current study, we apply a plethora of techniques, including NMR, fast mixing HDX-MS and nano electrospray ionisation ion mobility mass spectrometry (nano ESI-IM-MS) to study the differences in the conformation of aggregation prone and non-aggregation prone monomeric aSyn including phosphorylated aSyn, pS129, and familial aSyn mutants, A30P, E46K, H50Q, G51D, A53T and A53E. To investigate potential differences in residual structure and long-range interactions, we also performed experiments in the presence of calcium to purposefully skew the dynamic ensemble of conformations as calcium binds at the C-terminus and leads to charge neutralisation ^38^. Calcium has been shown to play a role in the physiological and pathological function of aSyn, as calcium binding at the C-terminus of aSyn facilitates interaction with synaptic vesicles and enhances aSyn aggregation rates ^8,39^. Furthermore, calcium-aSyn interaction is physiologically relevant as calcium buffering becomes dysregulated in PD and an increase in cytosolic calcium occurs ^40^. We therefore also investigated a panel of C-terminus mutants D115A, D119A and D121A which are within the calcium binding region^39^.

We conclude that the perturbation of the long-range interactions upon calcium binding to monomeric aSyn leads to an increase in N-terminal solvent exposure which correlates with aSyn’s aggregation propensity. The distribution of monomeric conformations of the aSyn ensemble is different between aSyn variants (e.g. familial mutants, pS129 aSyn) which populate these conformations to a different extent. The finding that different structural conformations can be identified as early as at the monomer level will be crucial in aiding the development of aSyn aggregation inhibitors stabilising native non-aggregation prone structures.

## Results

### pS129 and D121A have altered monomeric conformations compared to WT aSyn

We first investigated whether long-range interactions mediated by the C-terminus of aSyn were important in modulating monomer conformation and aggregation propensity of aSyn. We compared WT aSyn to post-translationally modified aSyn, phosphorylated at residue serine 129 (pS129) and to a mutant with reduced charge by mutating aspartate (D) to alanine (A) at residue 121 (D121A). This mutant was chosen from a panel of C-terminus D to A mutants, 115, 119 and 121, which reside in the region of divalent and trivalent cation binding sites ^8,41^. D121A aSyn was chosen for further investigation as it displayed a decreased aggregation rate using a thioflavin-T (ThT) based kinetic assay compared to D115A, D119A and WT aSyn. However, there was no difference in the overall structure as determined by fourier transform infrared spectroscopy (FTIR), which revealed disordered structures for all these variants, as discussed in SI (Figure S4-6).

We first compared the conformation of monomeric D121A and pS129 aSyn to WT aSyn using ^1^H-^15^N heteronuclear single quantum correlation (HSQC) spectra in solution. In comparison to WT aSyn there are chemical shift perturbations (CSPs) around the location of the D121A mutation, as expected, and some additional small CSPs at regions aar 1-10 and 70-80 (Figure 2a, green) which may indicate some disruption of long-range interactions in this mutant, as NMR measures an average of structural ensembles. CSPs in pS129 aSyn were present around the phosphorylated S129 residue compared to WT aSyn.

**Figure 2.**
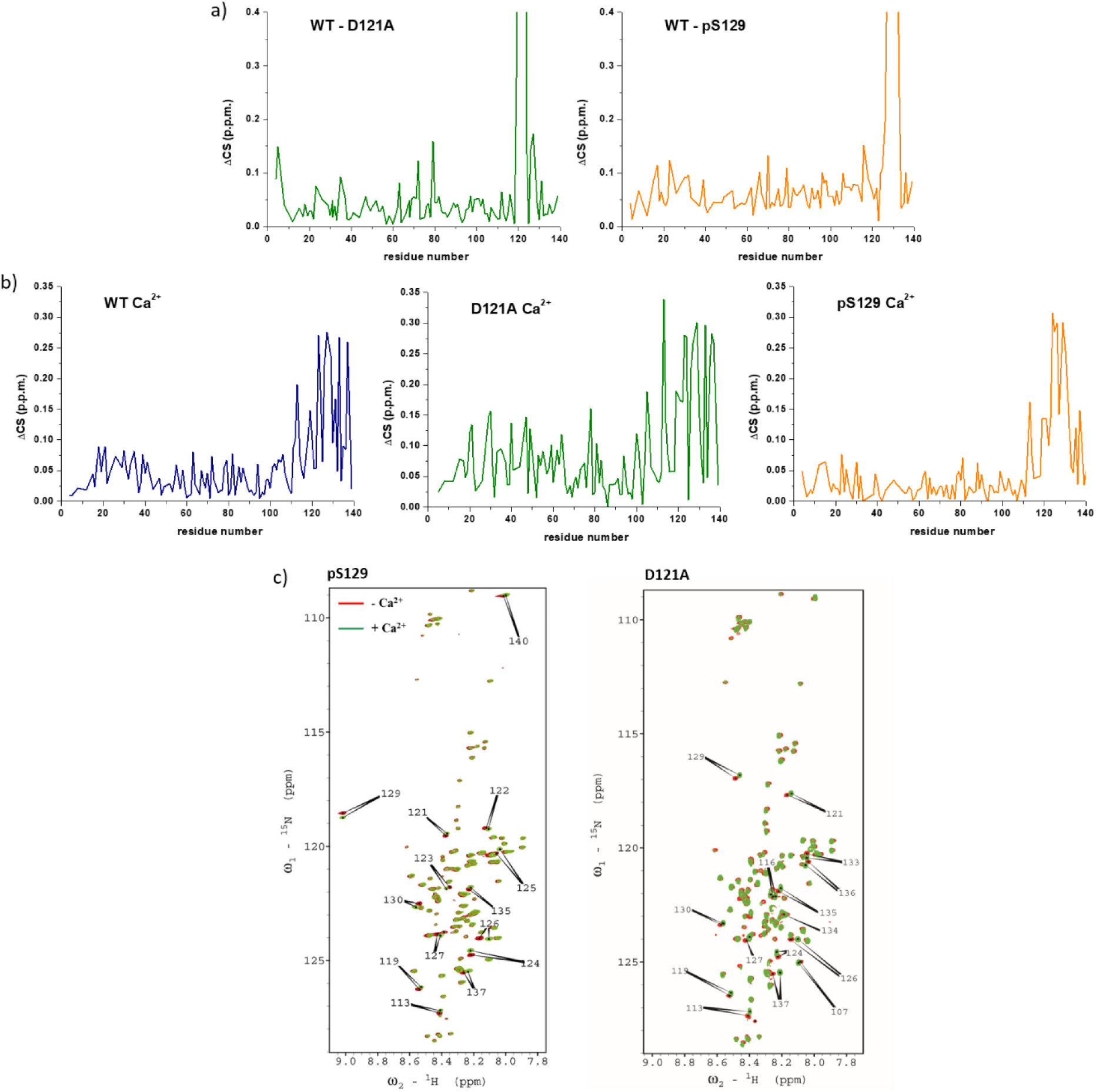
^1^H-^15^N spectra in the presence or absence of calcium of WT, D121A and pS129 aSyn indicate conformational changes between the different variants. (a) We compared chemical shift perturbations in the amide backbone of ^1^H-^15^N D121A aSyn to WT aSyn (green) and pS129 aSyn to WT aSyn (yellow) and observed CSPs around the location of the D to A mutation and the PO^4-^ of S129 at the C-terminus. Small CSPs were observed at residues 1-10 and 70-80 in D121A aSyn when compared to WT aSyn. (b) Chemical shift perturbations were observed at the C-terminus upon addition of 4.2 mM calcium for all, WT (blue), D121A (green) and pS129 (yellow) aSyn. Higher CSPs are observed for D121A compared to WT and pS129 aSyn. (c) 2D ^1^H-^15^N HSQC NMR spectra of pS129 and D121A aSyn in the absence (red) and in the presence of calcium (green). Major chemical shift perturbations in the presence of calcium are located at the C terminus (arrows with assigned amino acid residues) in both pS129 and D121A aSyn.

As CSPs of D121A and pS129 aSyn compared to WT aSyn were minimal aside the mutation/phosphorylation sites, we next investigated whether addition of calcium could skew the conformational ensemble of monomeric aSyn. Binding of calcium alters electrostatic interactions as it neutralises the negative charge at the C-terminus of aSyn ^8^. We observed significant CSPs at the C-terminus for all three aSyn samples, as shown previously for WT aSyn (Figure 2b) ^39^. For D121A aSyn, in addition to the main CSPs at the calcium binding area, we observe higher CSPs across the whole sequence when compared with WT and pS129 aSyn, which is unexpected since a point mutation usually only alters a very localised region of the sequence in a disordered protein ^42^ (Figure 2b, green). Furthermore, we observed no broadening of the NAC region when calcium was bound (Figure 2c), such as present in WT aSyn ^39^, suggesting that interactions with the NAC region were altered in D121A aSyn. Our data indicate that pS129 aSyn may also have altered long range interactions in comparison to WT aSyn. Firstly, there appear to be more localised C-terminus CSPs upon calcium binding compared to D121A and WT (Figure 2b, yellow). Residues involved in metal binding have previously been shown to be altered upon phosphorylation ^41^. Secondly, as observed for D121A aSyn, there was no broadening of the NAC region when calcium was bound (Figure 2c). These alterations in CSPs upon calcium binding suggest that long range interactions of pS129 and D121A aSyn are already altered compared to WT aSyn which may have implications in terms of the aggregation propensity of these mutants both in the absence and presence of calcium.

In order to determine whether the above structural changes we observed in the presence of calcium were simply due to changes in the affinity for calcium of the different aSyn variants, we performed calcium titrations experiments. We applied two different fitting algorithms to determine the dissociation constant. Using a previously applied model ^43^, but this time using a fixed ligand (calcium) number of 3 (which relates to the number of calcium ions bound as determined by MS with the same aSyn to calcium ratio), we obtained a *K_D_* of 95 (± 16) μM, 91 (± 16) μM and 69 (± 8) μM for WT, pS129 and D121A aSyn, respectively. Using the Hill equation, which also takes into account the level of cooperativity, we obtain a *K_D_* of 670 (± 51) μM, 670 (± 33) μM and 460 (± 27) μM for WT, pS129 and D121A aSyn, respectively. For all fittings, we get an *n* >1, indicating that calcium binds cooperatively to aSyn (Figure S7).

We next performed ThT-based kinetic assays to investigate whether the above observed conformational differences or calcium binding capacities of aSyn influenced the aggregation propensity of the three aSyn variants. The results of the assay showed that although D121A and pS129 aSyn had different charges at the C-terminus, both had a lower aggregation propensity than WT aSyn, particularly in the absence of calcium (Figure S8). The presence of calcium did increase the aggregation rate of D121A and pS129 aSyn in comparison to aggregation rates without calcium, however not to the same extent as the rate of WT aSyn. This was also reflected in the concentration of remaining monomer determined by size-exclusion chromatography high-pressure liquid chromatography (SEC-HPLC) (Figure S8 and S9). It thus appears that the affinity of aSyn to calcium does not influence aggregation rate per se as the affinity of D121A aSyn for calcium is higher than of WT and pS129 aSyn.

### D121A and pS129 aSyn are less exposed at the N-terminus compared to WT aSyn upon calcium addition

We further investigated whether the perturbations observed by NMR led to shielding or exposure of D121A and pS129 aSyn monomers using HDX-MS. This technique probes the submolecular structure and dynamics of proteins by employing hydrogen-deuterium exchange, and thus permit the identification of protein sequences that are more exposed to the solvent and/or less strongly hydrogen-bonded (deprotected). Binary comparison of the deuterium uptake profile between WT and D121A aSyn and WT and pS129 aSyn showed that both variants are not significantly different to WT aSyn (Figure 3a and b).

**Figure 3.**
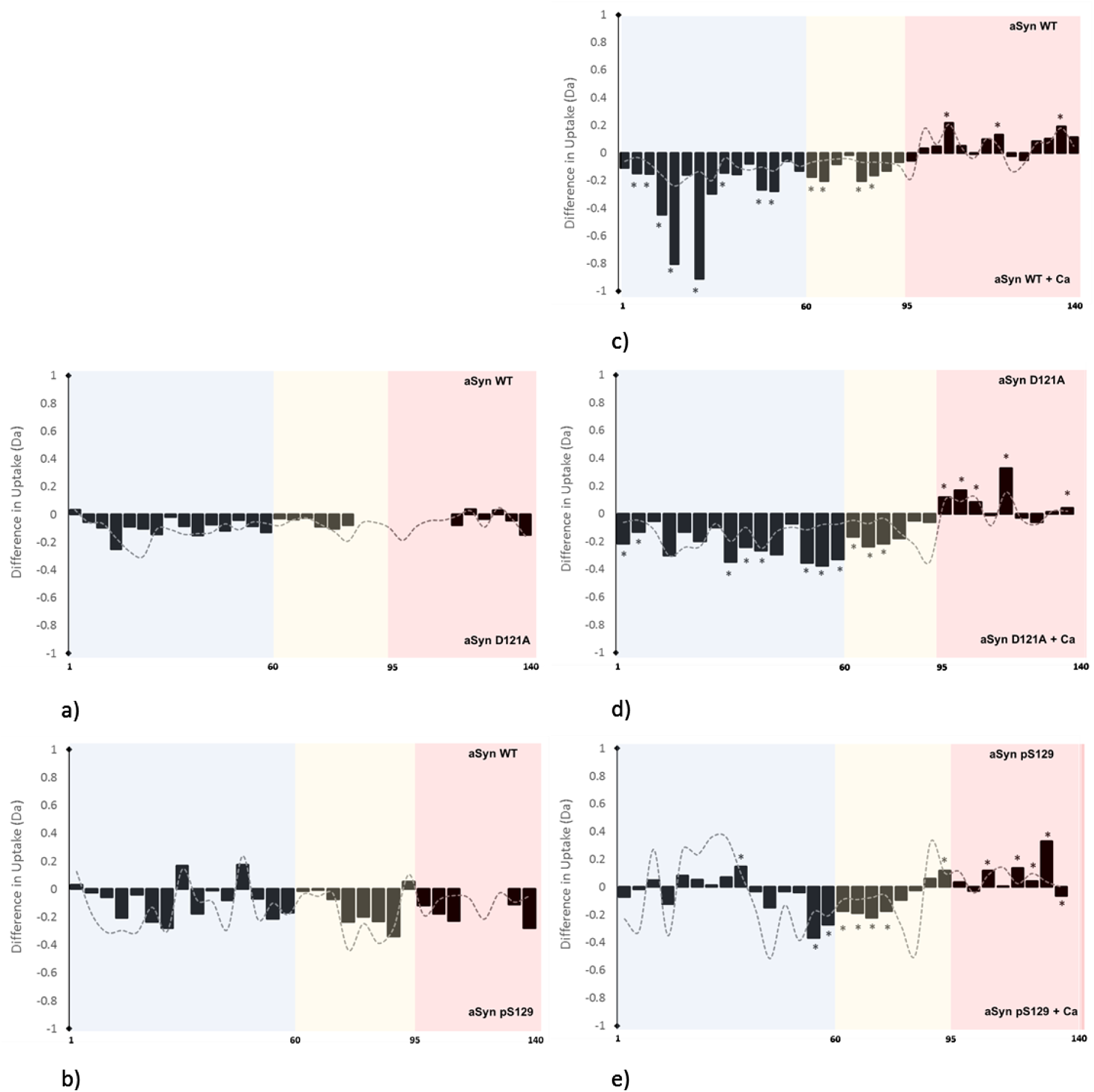
HDX-MS reveals different structural conformations in monomeric D121A and pS129 compared to WT aSyn. Bars represent differences in deuterium uptake along the sequence of differently compared aSyn variants (e.g. WT vs D121A aSyn) with the N-terminus labelled in blue, the NAC region in yellow, and the C-terminus of aSyn in red. Negative values represent increased deuterium uptake in the mutant (a,b) or in the calcium bound state (c-e), more solvent exposure, and less hydrogen bonding. Peptides containing the mutation were not comparable to WT aSyn and were removed from the data set, indicated by blank regions. Comparison of the deuterium uptake (in Dalton-Da) between (a) WT and D121A aSyn and (b) WT and pS129 aSyn showed that both were not significantly different to WT aSyn. (c) In the presence of calcium, WT aSyn becomes significantly more deprotected (more solvent exposed/less hydrogen bonded) at the N-terminus and the NAC region, and at the same time becomes solvent protected at the C-terminus. (d) D121A aSyn is significantly more deprotected at the N-terminus and the NAC region upon calcium addition and solvent protected at the C-terminus. (e) pS129 aSyn is deprotected at the NAC region upon calcium addition and solvent protected at the C-terminus. The grey trace signifies the error (1 s.d.) of six replicates collected per condition. Data acquired at each peptide were subjected to a Student’s t-test with a p-value ≤ 0.05 and significant differences are presented by a *.

We again used calcium to perturb the ensemble of conformations to compare alterations of long-range interactions between the three aSyn variants. Binary comparison of the deuterium uptake profile of monomeric WT aSyn revealed solvent protection at the C-terminus and significant deprotection at the NAC and the N-terminus region of aSyn in the presence of calcium compared to the absence of calcium (Figure 3c). This indicates that, when calcium is bound to aSyn, there is reduced exposure to the solvent or increased hydrogen bonding at the C-terminus of aSyn, where calcium binds, and deprotection of the N-terminus, as observed by CSPs using NMR. A similar behaviour was also observed for D121A and pS129 aSyn as, upon calcium binding, solvent protection is observed at the C-terminus and deprotection at the NAC region of aSyn (Figures 3d,e). However, while D121A aSyn has a more solvent exposed N-terminus, pS129 aSyn displays little difference in protection levels at the N-terminus. Despite both of these averaging structures being deprotected upon the addition of calcium, they are still more protected/less exposed compared to WT aSyn in the calcium-bound state (Figure 3c-e). Both NMR and HDX-MS indicate that D121A and pS129 aSyn have different ensembles of conformations compared to WT aSyn. Upon perturbation of the conformational ensemble by calcium binding, the degree of exposure of the N-terminus is much less pronounced in D121A and pS129 aSyn than in WT aSyn. This correlates with the reduced aggregation propensity in the calcium-bound state of D121A and pS129 aSyn, as determined by the ThT fluorescence kinetic assay.

### Familial aSyn mutants display differences in aggregation propensity and N-terminus solvent exposure compared to WT aSyn

To further investigate whether differences in the sub-molecular structure are apparent in the familial aSyn mutants A30P, E46K, A53T, A53E, H50Q, and G51D and whether this can influence aggregation rates, we first studied their aggregation kinetics in the presence and absence of calcium using ThT-based kinetic assays. Comparison of the fibrillisation rates of the familial mutants with WT aSyn in the absence of calcium shows that the aSyn mutants E46K, A53T, and H50Q aggregate faster than WT aSyn while the familial aSyn mutants A30P, A53E, and G51D aggregate more slowly than WT aSyn (Figure S10). Upon the addition of calcium, WT aSyn nucleation and elongation is enhanced, as previously shown ^39^, and for the fast aggregating aSyn mutants, A53T, E46K, and H50Q the aggregation rate is also enhanced upon the addition of calcium, similarly to WT aSyn. However, the slow aggregating aSyn mutants A30P, A53E, and G51D are either insensitive to calcium addition or aggregate more slowly. In order to determine whether the difference in aggregation was due to structural differences of the monomer, we again employed FTIR spectroscopy. The monomeric familial aSyn mutants all had an increased beta-sheet content, particularly A30P, E46K and A53T aSyn which were significantly different (Figure S11).

To gain more detailed and localised structural information than obtained by FTIR, we employed HDX-MS. The familial aSyn mutants A53T and A53E were selected from the familial mutants’ panel for this analysis, as they displayed different aggregation behaviour, even though the point mutations were localised on the same residue. Binary comparison of WT and A53T aSyn in the absence of calcium showed that there was no significant difference in deuterium uptake between the two protein states (Figure 4a), as observed for comparisons between WT and A53E aSyn (Figure 4b), and also A53T and A53E aSyn in the absence of calcium (Figure 4c). Upon the addition of calcium, solvent protection is observed at the C-terminus of A53T and A53E aSyn similar to WT and the other aSyn variants (see D121A and pS129 aSyn), indicating that the protection at this region is primarily due to calcium binding. At the same time, A53T aSyn is significantly deprotected at the N-terminus, at a similar level to WT aSyn, indicating a breaking of hydrogen bonding or a local unfolding event (Figure 4d,e). In contrast to the latter, for A53E aSyn (Figure 4f) no significant differences can be identified, neither at the NAC region nor at the N-terminus, upon calcium binding. Thus, the extent to which structural dynamics are perturbed in the N-terminal region in response to binding of calcium at the C-terminus correlates with an increase in the aggregation propensity of aSyn and its variants.

**Figure 4.**
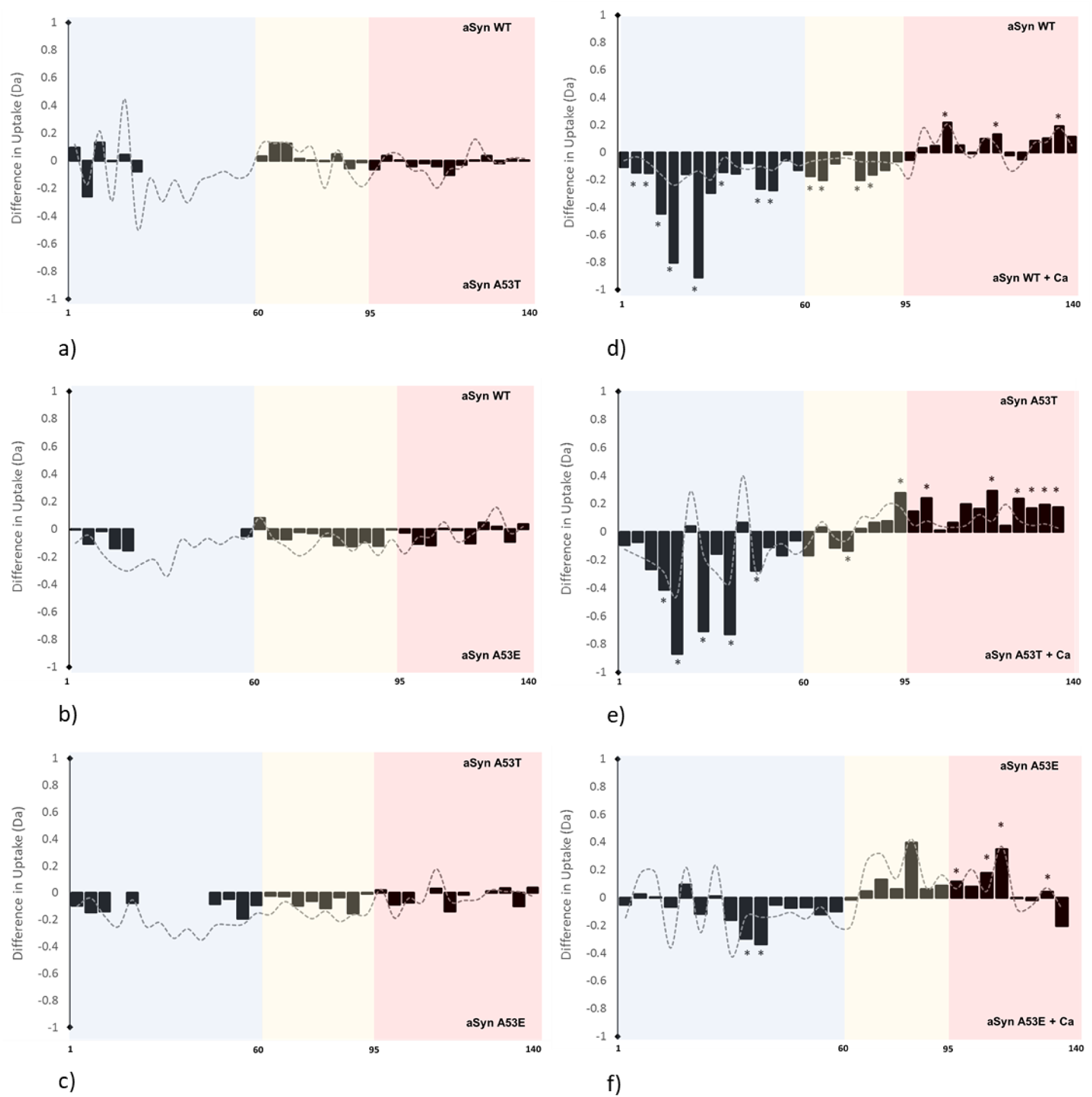
HDX-MS reveals different structural conformations of monomeric A53E compared to WT and A53T aSyn. Bars represent differences in deuterium uptake along the sequence of differently compared aSyn variants with the N-terminus labelled in blue, the NAC region in yellow, and the C-terminus in red. Negative values represent increased deuterium uptake in the mutant (a-c) or in the calcium bound state (d-f), more solvent exposure, and less hydrogen bonding. Peptides containing the mutation were not comparable to WT aSyn and were removed from the data set, indicated by blank regions. Difference in deuterium uptake (Da) between (a) WT and A53T aSyn, (b) WT and A53E aSyn and (c) A53T and A53E aSyn showed no significant differences throughout the sequence. In the presence of calcium, as shown in Figure 2c, (d) WT aSyn becomes significantly deprotected at the N-terminus and the NAC region, and more solvent protected at the C-terminus. (e) Similarly, A53T aSyn is significantly deprotected at the N-terminus and the NAC region upon calcium addition, and at the same time becomes solvent protected at the C-terminus. (f) A53E aSyn also becomes solvent protected at the C-terminus upon calcium addition but no significant changes are observed at the NAC region and most of the N-terminus. The grey trace signifies the error (1 s.d.) of six replicates collected per condition. Data acquired were subjected to a Student’s t-test with a p-value ≤ 0.05 and significant differences are presented by a *.

**Figure 4.**
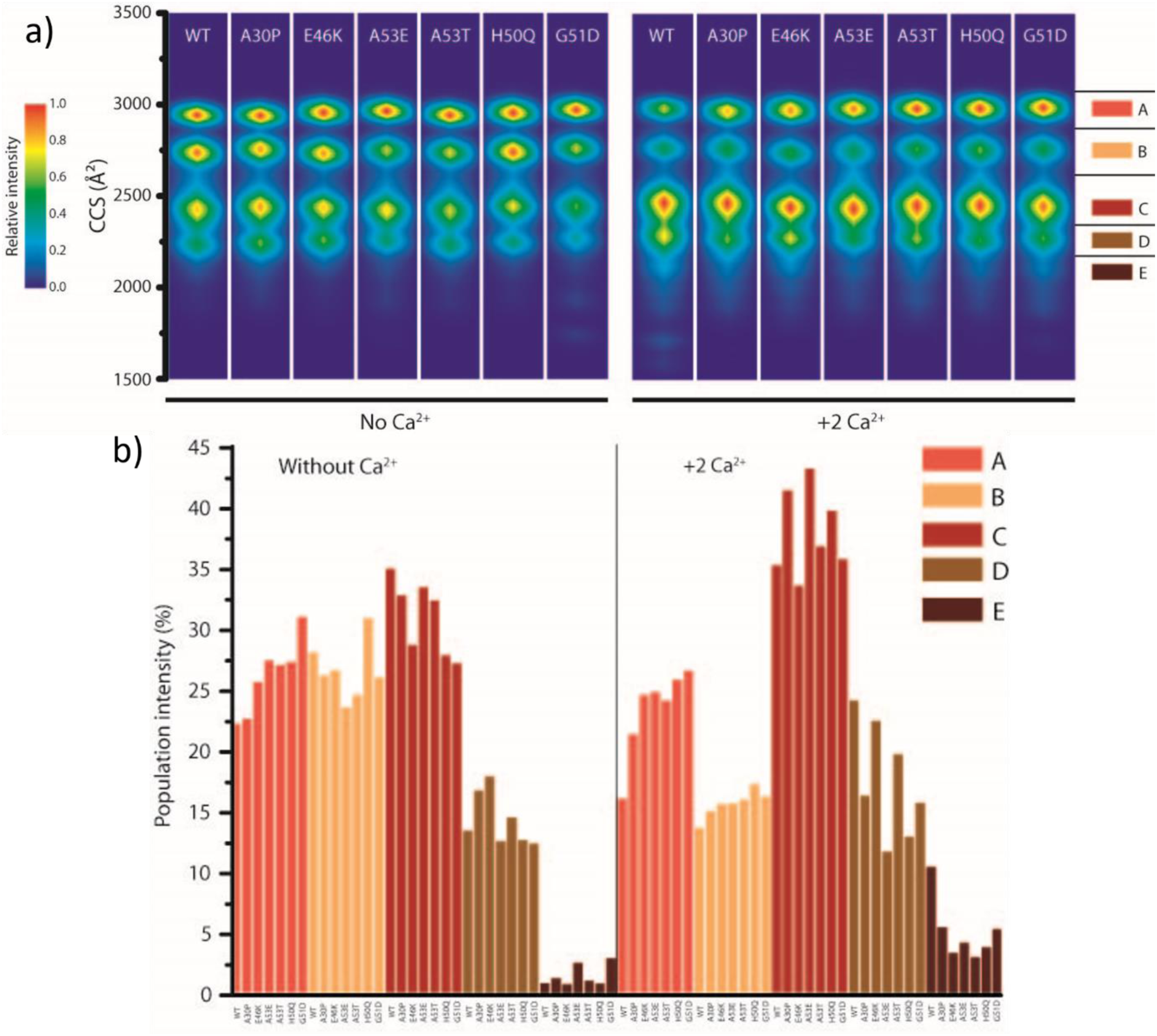
Nano-ESI-IM-MS reveals differences in aSyn conformation equilibria and compactions upon addition of calcium. Heat maps of aSyn conformations detected for the 8+ charge state based on intensity for (a) WT aSyn and the aSyn familial mutations, A30P, E46K, A53E, A53T, H50Q and G51D in the absence (No Ca^2+^) and presence of calcium at a 1:20 protein to calcium ratio, representing a two Ca^2+^ bound state (+2 Ca^2+^). Red represents the most populated CCS values by intensity. Upon addition of calcium, a higher proportion of aSyn have a lower collisional cross section (CCS) value (seen in the most populated region C), representing compaction. The area is separated into regions A, B, C, D, E at the right of the heat maps to denote and quantify the population of different conformations. (b) The conformational distributions of WT and mutant aSyn expressed as percentage were calculated from the area under each section or peak in the ATD and are presented in the absence (Without Ca^2+^) and presence of calcium (+2 Ca^2+^) and also in Tables S1, S2 including statistical analysis.

### The distribution of aSyn conformations is altered upon calcium binding

Techniques such as NMR and HDX-MS measure an ensemble of all conformations in the sample. Native nano-ESI IM-MS can instead characterise heterogeneous structural ensembles, which permits us to determine whether there is a difference in the distribution of conformations in the protein ensemble between different aSyn variants. The ion mobility of a protein is measured inside the mass spectrometer and determined by the number of collisions the protein ions have with gas molecules, influencing the drift (or arrival) time which is a direct correlation to the rotationally averaged size and shape of the particle. The larger the collision cross section (CCS), the more extended the conformation of aSyn, and the lower the CCS, the more compact the conformation. In the gas-phase, WT aSyn was found to have four main conformational distributions at the 8+ charge state (Figure 5a) which have been observed previously ^44–46^. A shoulder on the D peak of the protein profile in the arrival time distribution (ATD) suggests that there is an additional weak conformation present (E) at this charge state (Figure S12). As the charge state increases, the aSyn conformations become more extended due to a higher coulombic repulsion (Figure S13). The familial mutations also display four main co-existing conformational distributions, and no significant differences in overall compactness are observed compared to WT aSyn (Figure 5a, Figure S12). The ATD containing the main four CCS values at 8+ was subdivided into regions labelled A-D, with region E containing the least abundant and most compact conformation. Analysis of the data by both intensity (Figure 5a) and calculation of the area under each peak in the ATD, displayed as a percentage of the total population (Figure 5b) reveals no significant differences in the distribution of the ensemble of conformations between WT and mutant aSyn. There are however slight differences in the distribution of conformations between the different aSyn variants (Figure 5, Table S1 and S2).

**Figure 5.**
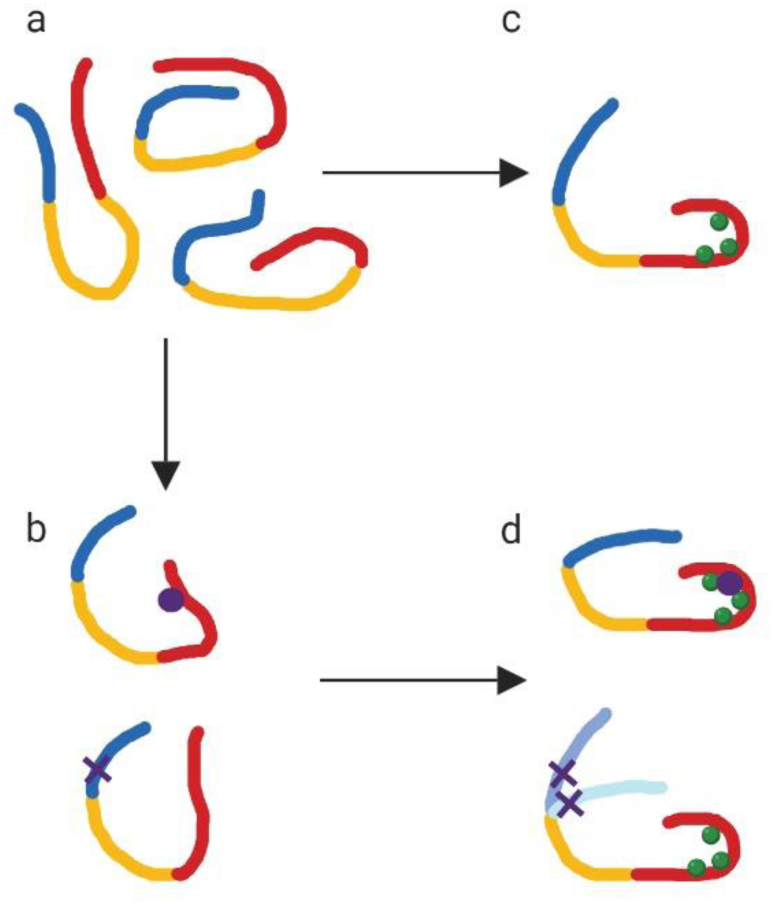
Cartoon representation of the effect of phosphorylation, mutation and calcium binding on aSyn. (a) aSyn is a dynamic ensemble of conformations in solution. Its solubility is maintained by long-range interactions between the N-terminus (blue), NAC region (yellow) and the C-terminus (blue) of aSyn. (b) Addition of a phosphate group to S129 (purple circle) or mutation (purple cross) alters the long-range interactions and skews the dynamic ensemble to favour or disfavour aggregation prone structures. (c) Addition of divalent cations such as calcium (green) leads to charge shielding, C-terminus collapse and exposure of the N-terminus of aSyn. (d) Calcium binding to phosphorylated or mutated aSyn leads to different perturbations of long-range interactions. Phosphorylation at S129 leads to a slightly different calcium binding region and to reduced solvent exposure of the N-terminus compared to WT aSyn. The aSyn mutation A53T (mid blue with purple cross) leads to N-terminus solvent exposure upon calcium binding at the C-terminus, similar to WT aSyn, however the aSyn mutation A53E (light blue with purple cross) leads to less N-terminus solvent exposure upon calcium binding.

Upon addition of calcium, compacted conformations are favoured in all aSyn variants, particularly conformation C, as indicated by more populated low CCS values (Figure 5a, +2Ca^2+^). Furthermore, analysis of the mass spectra showed that there were no significant differences in the distribution and maximum number of calcium ions bound to any of the aSyn variants, with, on average, two to three Ca^2+^ ions bound at a 1:10 protein to calcium ratio (Figure S14, S15).

## Discussion

As an intrinsically disordered protein, aSyn is constantly sampling a large conformational space and exists as an ensemble of conformations. The distribution of these conformations is significantly influenced by both changes in the sequence of the protein (e.g. mutations or PTMs) and the surrounding environment. Some of these structural conformers are expected to follow different aggregation pathways, possibly resulting in different fibril polymorphs and even leading to different disease outcomes. By understanding how genetic and environmental factors, such as mutations, PTMs and calcium, influence the dynamics of conformation and favour monomeric aggregation-prone structures we may begin to understand the initiation of misfolding pathways leading to different synucleinopathies, and subsequently how to disrupt them. However, here within lies the difficulty as the conformations monomeric aSyn samples are similar in size, charge and structure and are thus hard to detect using ensemble measurement techniques. Furthermore, it is difficult to discern differences in overall conformations when comparing mutant to WT aSyn. In this study, we used the biologically relevant ion, calcium, to perturb the conformational ensemble of aSyn structures and compared the differences in CSPs, hydrogen bonding/solvent exposure and distribution of conformations between aSyn and its mutants. In general, calcium has been shown to enhance the aggregation rate of aSyn ^39,47^, likely by a similar mechanism as low pH, whereby the reduction of the negative charge at the C-terminus leads to C-terminal collapse, altered long-range electrostatic interactions and enhanced hydrophobic interaction between the C-terminus and the NAC region which drives aggregation ^22,48–51^. We observed mutant specific differences in long-range interactions, compaction and solvent exposure compared to WT aSyn which will be discussed individually.

### Mutation and phosphorylation at the C-terminus alter long-range interactions

We first investigated the role of charge and long range interactions at the C-terminus by comparing two aSyn variants with reduced (D121A) and added (pS129) negative charge. We observed that both, D121A and pS129 aSyn displayed reduced aggregation rates in comparison to WT aSyn, indicating that increasing or decreasing charge by one residue at the C-terminus does not decrease or increase aggregation rates, respectively, as may have been expected. Many studies investigating the effect of charge on aSyn aggregation rates use C-terminus truncations which heavily disrupt and remove long-range interactions. aSyn truncated at Y133 and D135 leads to increased aggregation rates, but phosphorylation at S129 of these truncated aSyn reduces the aggregation rate, suggesting phosphorylation at S129 does have an inhibitory effect on aggregation^52^. Furthermore, calcium binding affinity does not correlate with aggregation rates, suggesting that altered residual monomeric structure at specific regions and not calcium binding or alteration of single charges per se influences aggregation rates. We observed only small differences in the conformational properties of the monomeric aSyn, as revealed by changes in CSPs and solvent exposure for D121A or pS129 compared to WT aSyn in the absence of calcium. This is likely due to the vast array of conformations sampled by the protein leading to minimal structural changes using protein ensemble measurement techniques such as NMR and HDX-MS. However, by perturbing the ensemble of conformations, as seen upon the addition of calcium, we could observe differences in CSPs and deuterium uptake upon calcium binding. pS129 aSyn has a different calcium-binding region and displays no broadening in the NAC region compared to WT aSyn, while D121A has a higher degree of CSPs across the sequence, but also no NAC region broadening, suggesting long-range interactions with the NAC region in both these mutants are altered before calcium binds. Broadening in the NAC region has been observed for WT aSyn when bound to calcium ^39^ and at low pH, suggesting enhanced interactions the between calcium binding region, namely the charge neutralised C-terminus, and the NAC region ^22^. Regarding aSyn’s propensity to aggregate, we observe, using HDX-MS, that the N-terminus of pS129 aSyn is not solvent exposed upon calcium binding, whereas for D121A aSyn, although the N-terminus is slightly exposed, is significantly less exposed compared to WT aSyn upon calcium binding, correlating to the decreased aggregation propensity of D121A and pS129 aSyn. Previous HDX-MS studies have suggested that the N-terminus of aSyn may be involved in aggregation as there is heterogeneity in its solvent exposure during aggregation which is also linked to fibril morphology, while the C-terminus remains completely solvent exposed ^53–55^. Here, we show that the extent of N-terminus (aar 1-60) and NAC region (aar 61-95) exposure, based on the amount of deuterium exchange which is determined by ease of accessibility, is correlated to the aggregation propensity of aSyn, where WT>D121A>pS129 in terms of N-terminus exposure, but the opposite is observed for aggregation propensity. Importantly, we *only* observe N-terminus exposure upon the disruption of long-range interactions with the C-terminus upon calcium binding. The C-terminus thus greatly influences aggregation propensity, as it is important in maintaining long range interactions and solubility of monomeric aSyn ^56,57^. Mutation of proline^58^, or glutamate^57^ residues to alanine at the C-terminus increases aSyn aggregation propensity, yet mutation of tyrosine^27^ residues to alanine decreases aSyn aggregation propensity. This suggests that mutations of specific residues alter specific long-range interactions which subsequently influence the distributions of conformations within the dynamic ensemble and thus the aggregation propensity of aSyn and its variants. Even phosphorylation of S129 compared to Y125 aSyn (only 4 aar apart) leads to differences in aggregation rates, as pY125 does not influence the aggregation rate compared to WT aSyn ^59^, neither the binding capacity of aSyn to nanobodies nor the alteration of metal binding sites, indicating very site-specific functions and interactions are present at the C-terminus ^41,59^. Although D121A aSyn is not a naturally occurring mutation, by comparing it to pS129, we can explore how altering long-range interactions of monomeric aSyn at the C-terminus by mutation, PTM or addition of divalent cations leads to altered interactions with the NAC region and different levels of solvent exposure at the N-terminus. Mutations leading to altered long-range interactions lead to a favouring of non-aggregation prone conformations in pS129 and D121 aSyn presented here, but under other environmental conditions not tested here, a favouring of aggregation prone conformations may also occur, and thus both sequence and environment are important in determining aSyn’s aggregation propensity.

### Familial aSyn mutants display different long-range interactions

While all familial aSyn point mutations reside at the N-terminus, the putative calcium binding site is at the C-terminus ^60^. Certainly, the presence of a familial aSyn mutation has been shown to alter interactions between the mutation site and the C-terminus by NMR compared to WT aSyn, with regional differences apparent for each aSyn mutant in physiological and mildly acidic conditions ^23,35^. In the present study, we observe that the familial aSyn mutants have different aggregation kinetics in response to calcium, which is understandable considering the mutation regions interact with the calcium binding regions at the C-terminus. Using HDX-MS, we observe a difference in the level of solvent protection at the N-terminus of A53T and A53E aSyn upon binding of calcium suggesting that indeed different long-range interactions are present. A local unfolding event (deprotection) at the N-terminus and the NAC region of WT and A53T aSyn correlates with their increased aggregation propensity upon calcium addition. No such deprotection was observed for the aSyn mutant A53E, whose aggregation kinetics were similar in the presence and absence of calcium. We expect that the release of long-range contacts with the C-terminus leads to exposure of the N-terminus and the NAC region, which is observed in WT and A53T aSyn, to be a major factor influencing aSyn aggregation kinetics. Several experiments have shown the importance of the N-terminus in modulating aSyn aggregation. The presence of repeating units, KTKE, across the N-terminus seems to have an important role because addition, deletion, swapping and their spacing lead to differential aggregation rates ^61,62^. Furthermore, cross linking G41C and V48C leads to inhibition of aggregation ^63^, and targeting proteins that bind to aar 37-54 prevents aggregation ^64,65^. It must be noted that all the familial aSyn mutations are also present at the N-terminus and several studies have shown the mutant aSyn have destabilised monomeric structures and altered long-range interactions compared to WT aSyn ^66–70^. The altered stability combined with a lower propensity to form an α-helix, as observed by FTIR, at the N-terminus compared to WT aSyn, may skew the distribution of the conformational ensemble leading to conformations with high propensity to form oligomers and fibrils ^67,68,71,72^. We observe increased N-terminus solvent exposure for the ‘fast aggregating’ mutant A53T compared to the ‘slow aggregating’ mutant A53E. In another study, the ‘fast aggregating’ mutant, E46K, also displays an increased solvent exposure across its sequence, and, in addition, N-terminus residues are involved in oligomerisation ^35^. The fact that such differences in aSyn long-range contacts and an increase in N-terminus solvent exposure can be identified as early as the monomer level, and correlate to the propensity to fibrillise is important, and a step towards understanding early events in the misfolding pathway and how structure and environmental factors may influence aggregation propensity of the monomer.

To gain insight into the mechanisms by which mutations associated with familial PD alter aggregation kinetics of aSyn, we sought to determine whether there were differences in the distribution of monomer conformations between the familial aSyn mutants and WT aSyn using native nano-ESI-IM-MS. We observed the presence of more than one population of conformations of aSyn in the gas phase, indicating that aSyn may also take on a multitude of different conformations in solution. Upon calcium addition, the conformational ensemble shifted towards favouring compacted structures in all aSyn mutants as the C-terminus collapses due to charge neutralisation upon binding to calcium, which has also been previously observed when Mn^2+^ and Co^2+^ bind WT aSyn ^44^. Divalent cation binding is specific to the C-terminus, leading to compaction of aSyn conformations, non-specific binding of monovalent ions (e.g. K^+^, Na^+^) leads to extended structures being favoured due to charge shielding of the N- and C-termini (A.D. Stephens, et. al., in preparation). Of note, the CCS values most favoured in the calcium-bound state in region C are also present in the non-calcium bound state, but to a lesser extent, it is thus possible that calcium binding leads to a bias towards structures that are already available to the monomer in the calcium-free state. The increase in aggregation propensity of WT aSyn upon calcium binding cannot only be explained by charge neutralisation at the C-terminus as all familial aSyn mutants should have responded in the same way to calcium which was not the case. It is more likely that the difference in aggregation propensity is a result of perturbed long-range interactions between the C-terminus, and thus the sequence and its interaction with the local environment, which skew the population towards more aggregation prone structures. Multiple conformations are likely present in each CCS value group, suggesting that nano-ESI-IM-MS today may not have the resolution to determine differences in the ensemble of conformations between WT and mutant aSyn.

## Conclusions

As an IDP, aSyn samples many different conformations making it difficult to identify specific aggregation prone conformations, particularly using ensemble measurement techniques. By comparing the submolecular structure of calcium-bound aSyn variants, we instead sampled a skewed population and inferred differences in structure and aggregation propensity as part of a response to calcium binding at the monomer level. We attribute the increase in aggregation propensity upon calcium binding to structural perturbation, as we observe no correlation in aSyn’s aggregation propensity to its affinity to calcium, the number of calcium ions bound or charge neutralisation at the C-terminus. Instead, we observe different responses to calcium based on the presence of different long-range interactions which are likely already altered in non-calcium bound forms of the familial mutants and upon the addition of PO^4-^ at S129 (Figure 5). Calcium binding leads to further disruption of intramolecular interactions with the C-terminus leading to unfolding and solvent exposure of the N-terminus.

The extent of N-terminus solvent exposure upon C-terminal binding of calcium correlates with the aggregation propensity of aSyn; pS129, A53E, and to some extent D121A aSyn were less solvent exposed at the N-terminus and had a reduced aggregation propensity compared to WT and A53T aSyn which were more aggregation prone and more solvent exposed at the N-terminus and at the beginning of the NAC region (Figure 5). It remains to be determined whether these different structures of monomeric aSyn can be isolated or whether they have different toxicity in cells. This would be an important step in rationalising the molecular mechanism of PD and other synucleinopathies, by identifying which local environmental factors bias the monomeric population towards the most disease-relevant conformers. Finally, targeting the most disease-relevant monomeric structures of aSyn and to convert them into ‘normal’ functional structures could open up a new chapter in the design of therapeutics against PD and other synucleinopathies.

## Methods

### Purification of aSyn

Human wild-type (WT) alpha-synuclein was expressed using plasmid pT7-7. ASyn mutations D115A, D119A, D121A, A30P, E46K, A53T, A53E, H50Q and G51D were introduced using the QuikChange Lighting Site-Directed Mutagenesis K (Agilent Technologies LDA UK Limited, UK) and confirmed by sequencing (Source Bioscience, Nottingham, UK). The plasmids were heat shocked into Escherichia coli One Shot^®^ BL21 STAR™ (DE3) (Invitrogen, Thermo Fisher Scientific, Cheshire, UK) and purified as previously described ^18^. Briefly, expressed aSyn was purified using ion exchange chromatography (IEX) on a HiPrep Q FF 16/10 anion exchange column (GE Healthcare, Uppsala, Sweden). aSyn was then further purified on a HiPrep Phenyl FF 16/10 (High Sub) hydrophobic interaction chromatography (HIC) column (GE Healthcare). aSyn was extensively dialysed against 20 mM Tris pH 7.2 and concentrated using 10 k MWCO amicon centrifugal filtration devices (Merck KGaA, Darmstadt, Germany) and stored at −80 °C until use. Before experiments 1 mL of aSyn was further purified using a Superdex 75 pg 10/300 GL size exclusion chromatography (SEC) column (GE Healthcare) to obtain monomeric protein. Purification was performed on an ÄKTA Pure (GE Healthcare). Protein concentration was determined by measuring the absorbance at 280 nm on a Nanovue spectrometer using the extinction coefficient 5960 M^-1^cm^-1^.

Protein purity was analysed using analytical reversed phase chromatography. Each purification batch was analysed using a Discovery BIO Wide Pore C18 column, 15cm x 4.6mm, 5µm, column with a guard cartridge (Supelco by Sigma-Aldrich) with a gradient of 95% to 5% H_2_O + 0.1% trifluroacetic acid (TFA) and acetonitrile + 0.1% TFA at a flow-rate of 1 mL/min. The elution profile was monitored by UV absorption at 220 nm and 280 nm on an Agilent 1260 Infinity HPLC system (Agilent Technologies LDA UK Limited, UK) equipped with an autosampler and a diode-array detector (a representative chromatograph is shown in Figure S1A). Protein purity fell between 89 - 96 %.

### Purification of aSyn for nuclear magnetic resonance experiments

E. coli was grown in isotope-enriched M9 minimal medium containing 15N ammonium chloride, and 13C-glucose similar to our previous protocol ^73^. Briefly, to isolate expressed aSyn the cell pellets were resuspended in lysis buffer (10mM Tris-HCl pH 8, 1mM EDTA and EDTA-free complete protease inhibitor cocktail tablets (Roche, Basel, Switzerland), 0.2 mM phenylmethylsulfonyl fluoride (PMSF) and Pepstatin A and lysed by sonication. The cell lysate was centrifuged at 22,000 g for 30 min to remove cell debris and the supernatant was then heated for 20 min at 90 °C to precipitate the heat-sensitive proteins and subsequently centrifuged at 22,000 g. Streptomycin sulfate (Sigma-Aldrich) 10mg/ml was added to the supernatant to precipitate DNA. The mixture was stirred for 15 min followed by centrifugation at 22,000 x g, then repeated. Ammonium sulfate 360 mg/ml was added to the supernatant precipitate the protein aSyn. The solution was stirred for 30 min and centrifuged again at 22,000 x g. The resulting pellet was resuspended in 25mM Tris-HCl, pH 7.7 and dialysed overnight. The protein was purified by IEX on a HiPrep Q FF anion exchange column (GE Healthcare) and then further purified by SEC on a HiLoad 16/60 Superdex 75 prep grade column (GE Healthcare). All the fractions containing the monomeric protein were pooled together and concentrated by using amicon 10 k MWCO centrifugal filtration devices (Merck).

### Purification of phosphorylated serine 129 aSyn

WT α-syn ^13^C/^15^N-labelled (α-syn ^13^C/^15^N) was expressed and purified as previously described ^71,74^. Briefly, *E.coli* BL21(DE3) cells were transfected with a pT7-7 plasmid containing WT aSyn and was cultured in an isotopically supplemented minimal media according to a previously described protocol ^71^, then aSyn ^13^C/^15^N was purified using an anion exchange chromatography followed by reversed-phase HPLC (RP-HPLC) purification using a Proto 300 C4 column and a gradient from 30 to 60% B over 35 min at 15 ml/min, where solvent A was 0.1% TFA in water and solvent B was 0.1% TFA in acetonitrile, the fractions containing the protein were pooled and lyophilized and the protein was stored at -20°C. For the preparation of phosphorylated S129 α-syn ^13^C/^15^N (aSyn ^13^C/^15^N pS129), we used a previously established protocol using PLK3 kinase to introduce selectively the phosphorylation at S129 ^75^. The WT aSyn ^13^C/^15^N was resuspended in the phosphorylation buffer (50 mM HEPES, 1 mM MgCl2, 1 mM EGTA, 1 mM DTT) at a concentration of ∼150 µM and then 2 mM of ATP and 0.42 μg of PLK3 kinase ( Invitrogen) per 500 μg of protein were added. The enzymatic reaction was left at 30°C overnight without shaking. Upon complete phosphorylation, as monitored by mass spectroscopy (LC/MS), aSyn ^13^C/^15^N pS129 was purified from the reaction mixture by RP-HPLC using an Inertsil WP300-C8 semiprep column. Finally, the fractions containing the protein of interest were pooled and quality control of aSyn ^13^C/^15^N pS129 was performed using mass spectroscopy, UPLC, and SDS-PAGE, the protein was 99.89% phosphorylated (Figure S1B-D).

### Solution nuclear magnetic resonance (NMR)

In order to probe the structure and thermodynamics of calcium binding with aSyn WT, pS129 and D121A at a residue specific level, we employed a series of ^1^H-^15^N HSQC experiments using different concentrations of Ca^2+^ (0.0 mM to 4.2 mM) and a fixed concentration of aSyn (200 μM). NMR experiments were carried out at 10°C on a Bruker spectrometer operating at ^1^H frequencies of 800 MHz equipped with triple resonance HCN cryo-probe. The ^1^H-^15^N HSQC experiments were recorded using a data matrix consisting of 2048 (t_2_, ^1^H) × 220 (t_1_, ^15^N) complex points. Assignments of the resonances in ^1^H-^15^N-HSQC spectra of aSyn were derived from our previous studies.

The perturbation of the ^1^H-^15^N HSQC resonances was analysed using a weighting function:

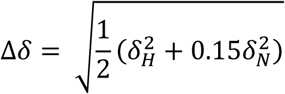

The titration enabled calculating the fraction of bound aSyn, *χ*_*B*_, as a function of [Ca^2+^].

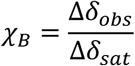

Where the Δδ_*obs*_ is the chemical shift perturbation of the amide group of a residue of aSyn at a given [*Ca*^2+^] and Δδ_*Sat*_ is the perturbation obtained with calcium saturation. *χ*_*B*_ values were obtained as a function of [*Ca*^2+^] for every residue of the protein for which resolved peaks in the ^1^H-^15^N HSQC are available. A global *χ*_*B*_ was calculated by average the fractions corresponding to residues associated with major resonance perturbations in the presence of calcium.

In order to obtain the apparent dissociation constant we used two different models. As first, we employed a model based on previous investigations ^39^:

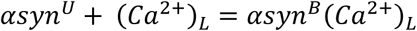

Where *αsyn*^*U*^ and *αsyn*^*B*^ indicate free and calcium bound aSyn, L indicates the number of Ca^2+^ interacting with one aSyn molecule, and the overall concentration of aSyn in this equilibrium is given by

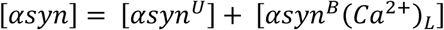

the apparent dissociation constant in this model corresponds to:

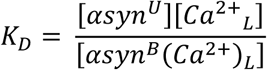

Which provides a formula for *χ*_*B*_:

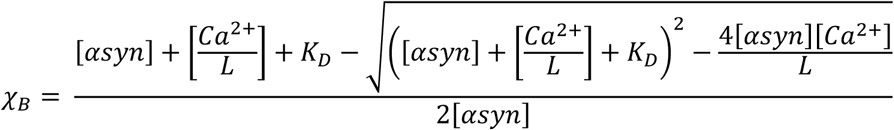

When using this fitting model in the case of WT aSyn we obtained a *K_D_* of 21 (± 5) μM and an L of 7.8 (± 0.51) ^39^. Based on the present MS data, we here fixed the value of L to 3.

We then used a different model that accounts for the cooperativity of the binding. In particular, we used the Hill equation to fit our data:

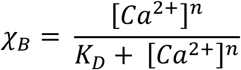

Where the Hill coefficient, *n*, describes the cooperativity of the binding. A *n* value higher than 1 indicates a positive cooperativity for the binding.

### Hydrogen-deuterium exchange mass spectrometry (HDX-MS)

Hydrogen exchange was performed using an HDX Manager (Waters, USA) equipped with a CTC PAL sample handling robot (LEAP Technologies, USA). Samples of aSyn in protonated aqueous buffer (20 mM Tris, pH 7.2) were diluted 20-fold into deuterated buffer (20 mM Tris, pD 7.2) at 20 °C, initiating hydrogen exchange. The same was performed for the calcium condition, in protonated and deuterated buffers containing 1 mM CaCl_2_ (20 mM Tris, pH 7.2, 1 mM CaCl_2_, and 20 mM Tris, pD 7.2, 1 mM CaCl_2_). The protein was incubated for 30 s in the deuterated buffer and six replicates were collected per condition. Hydrogen exchange was arrested by mixing 1:1 with pre-chilled quench buffer (100 mM Tris, 8 M Urea, pH 2.45 at 0 °C). The protein was then digested into peptides on a pepsin column (Enzymate, Waters) and the peptides were separated on a C18 column (1×100 mm ACQUITY UPLC BEH 1.7 μm, Waters) with a linear gradient of acetonitrile (3-40 %) supplemented with 0.2 % formic acid. Peptides were analysed with a Synapt G2-Si mass spectrometer (Waters). The mass spectrometer was calibrated with NaI calibrant in positive ion mode. Α clean blank injection was ran between samples to minimise carry-over. Peptide mapping of aSyn, where peptides were identified by MS/MS fragmentation, was performed prior to the hydrogen exchange experiments and analysed using ProteinLynx Global Server-PLGS (Waters). Peptide mapping of aSyn yielded coverage of 100% of aSyn with a high degree of redundancy (5.32) (Figure S2). The data pertaining to deuterium uptake (representative data presented in Figure S3) were analysed and visualised in DynamX 3.0 (Waters) and GraphPad Prism (GraphPad Software, US). No correction was made for back-exchange.

### Nano electrospray ionisation ion mobility mass spectrometry (Nano ESI-IM-MS)

A final concentration of 20 µM AS (WT or mutant) was obtained in 20 mM ammonium acetate (Sigma Aldrich, St. Louis, MO, USA) pH 7 and measured as a control. CaCl_2_ (Merck, Darmstadt, Germany) was dissolved in deionised H_2_O and added to the sample with a final concentration ranging between 200 µM and 400 µM. The samples were incubated for 10 minutes at room temperature before measuring. Nano-ESI (ion mobility-) mass spectrometry (nano-ESI-IM-MS) measurements were performed on a Synapt G2 HDMS (Waters, Manchester, U.K.) and analysed using Masslynx version 4.1 (Waters, Manchester, U.K.). For infusion into the mass spectrometer, home-made gold-coated borosilicate capillaries were used. The main instrumental settings were: capillary voltage 1.4-1.8 kV; sampling cone 25 V; extraction cone 1 V; trap CE 4 V; transfer CE 0 V; trap bias 40 V. Gas pressures used throughout the instrument were: source 1.5-2.7 mbar; trap cell 2.3 x 10-2 mbar; IM cell 3.0 mbar; transfer cell 2.5 x 10-2 mbar.

## Supplementary Data

**Figure S1.**
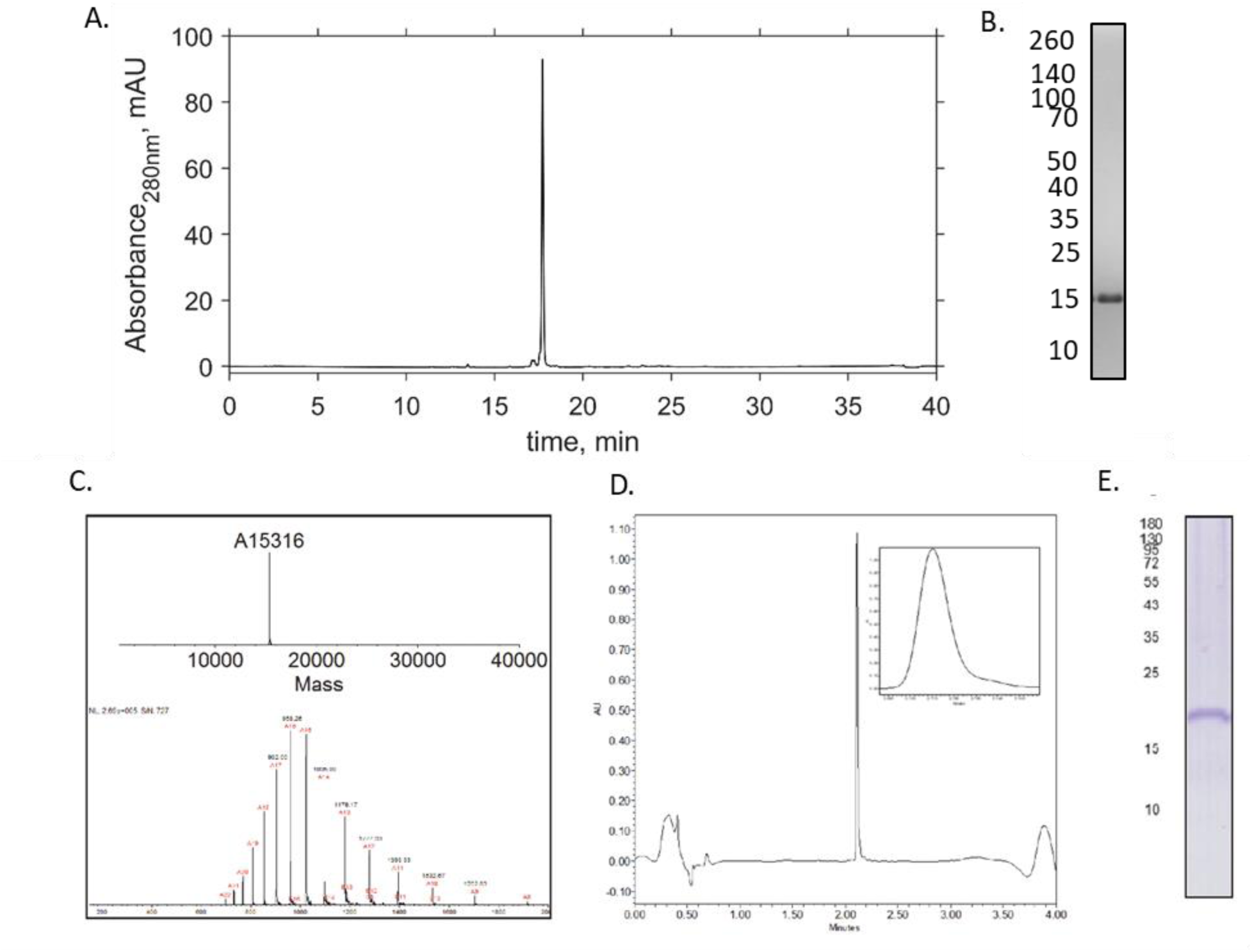
Analysis of WT and ^13^C/^15^N pS129 aSyn purity and percentage phosphorylation. (A) Representative chromatograph of WT aSyn analysed by analytical reverse phase (RP)-HPLC, 50 pL of aSyn was injected onto a Discovery Bio Wide Pore C18-5 column and eluted using a gradient of 5% acetonitrile + 0.1% acetic acid to 95% acetonitrile + 0.1% acetic acid with H_2_O + 0.1% acetic acid over 40 minutes at 1 ml/min. Percentage purity of aSyn was 93.9% based on absorbance at 280 nm. (B) Coomassie blue staining of SDS-PAGE gel of WT aSyn. (C) Phosphorylation of ^13^C/^15^N labelled aSyn at S129 was confirmed by mass spectrometry (expected mass 15.333 kDa). (D) Ultra performance liquid chromatography (UPLC) and (E) coomassie blue staining of SDS-PAGE gel was used to determine aSyn purity. The labelling percentage of S129 was 99.89%.

**Figure S2.**
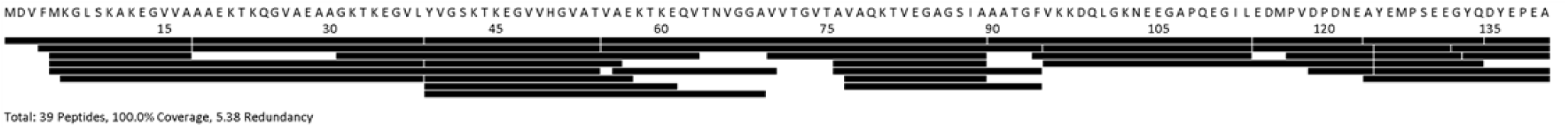
Peptide coverage map of aSyn using HDX-MS. Peptide mapping of aSyn was performed on a pepsin column prior to HDX-MS experiments, yielding 100% coverage of aSyn with a high degree (5.38) of redundancy. Peptides were identified by MS-MS fragmentation with ProteinLynx Global Server (Waters).

**Figure S3.**
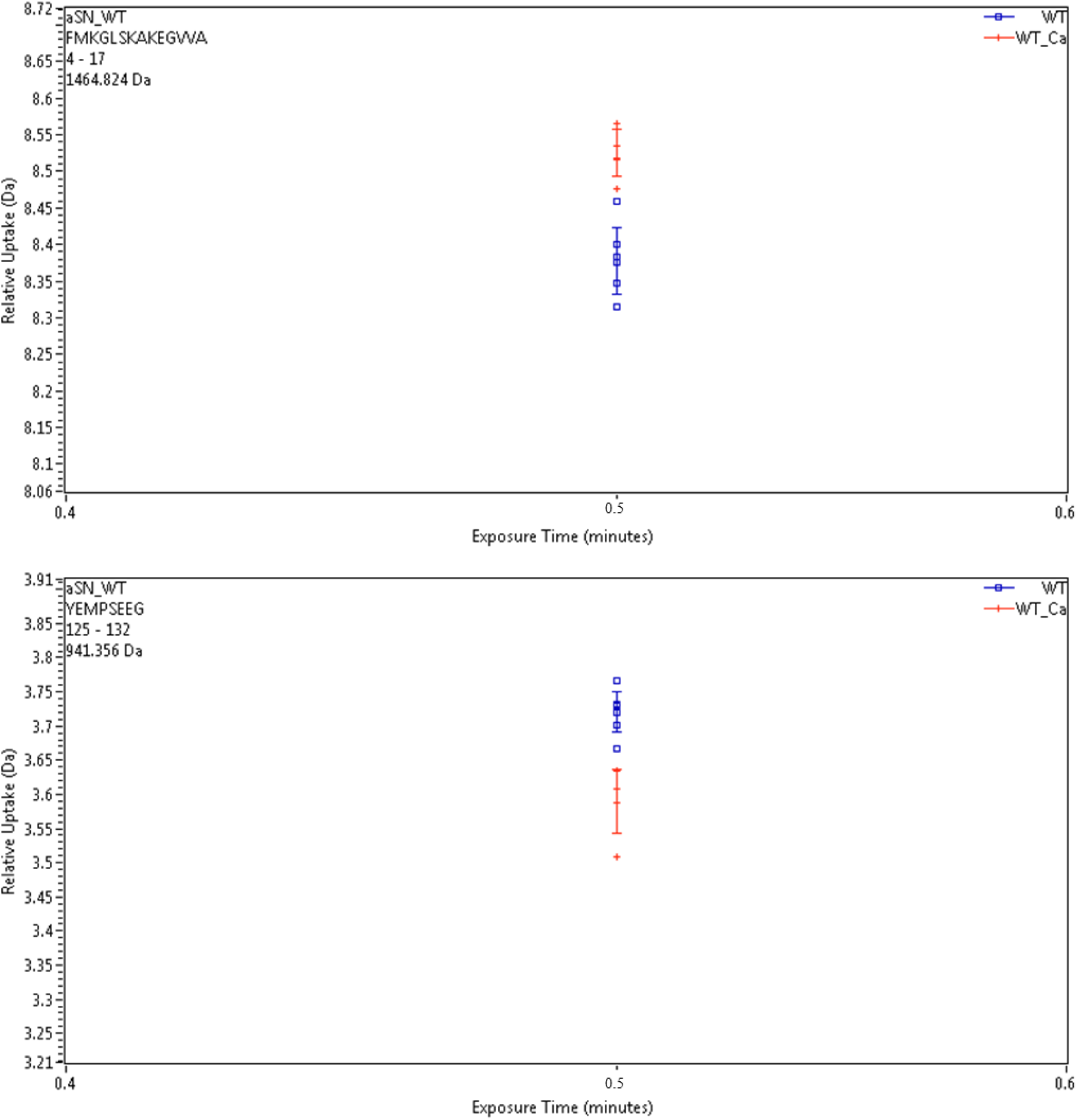
Deuterium uptake plots, measured in Dalton (Da), for representative peptides containing aar 4-17 (N-terminus) and aar 125-132 (C-terminus) of WT aSyn in the presence (red) and absence (blue) of calcium. Six replicates were collected per condition (+/-calcium) for a 30 s labelling timepoint. Error indicates 1 s.d. No correction was made for back-exchange. Uptake data for all the peptides were collected and further analysed to form Figures 3 and 4. Discussion of ThT assays and FTIR of D to A mutants Thioflavin T (ThT)-based kinetic assays showed little difference in aggregation rates for D115A, D119A, D121A and WT aSyn in the absence of calcium, but in the presence of calcium the D121A aSyn mutant had a significantly reduced aggregation rate compared to the other aSyn variants (Figure S2). Furthermore, D121A aSyn did not aggregate into fibrillary structures under these conditions as determined by AFM (Figure S3), suggesting that D121A may not bind calcium or had a different, non-aggregation prone monomeric structure. Analysis of the structure by FTIR showed that there were no significant differences in all D to A mutant structures compared to WT, which was overall disordered (Figure S4).

**Figure S4.**
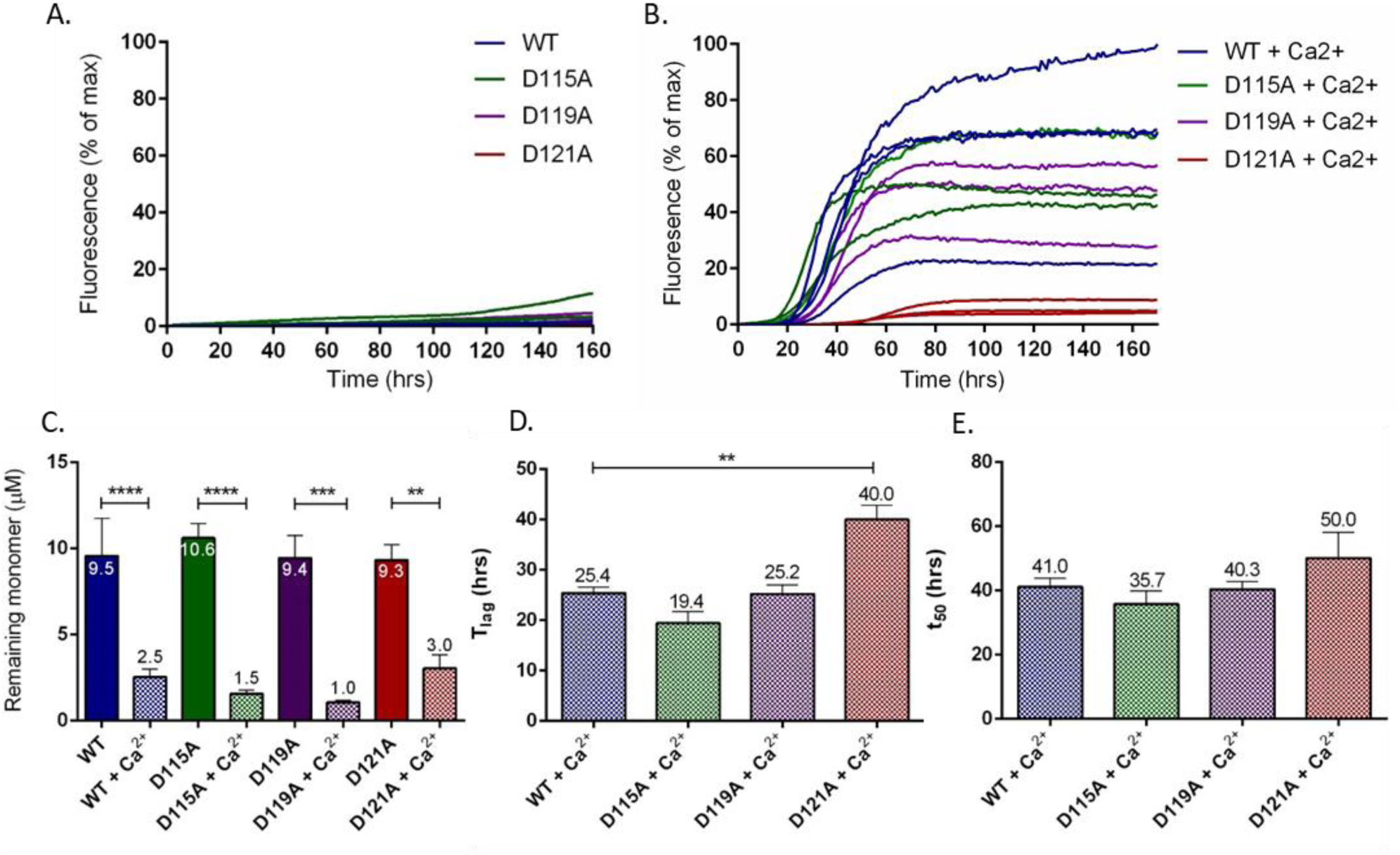
ThT-based aggregation assay reveals that D121A aSyn does not aggregate to the same extent as WT, D115A and D119A upon incubation with calcium. (A) The average of individual experiments is shown as ThT fluorescence intensity plotted as % of maximum fluorescence per plate of WT (blue), D115A (green), D119A (purple) and D121A aSyn (red) and (B) in the presence of 2.5 mM CaCl_2_. 50 μM aSyn was incubated with 10 μM ThT in a 384 well plate with orbital agitation at 300 rpm for 5 minutes before each read every hour for 160 hours. (C.) Remaining monomer was measured using SEC-HPLC, 35 μL of monomer from each well in the ThT assay were analysed on an AdvanceBio SEC 130Å column equilibrated in 20 mM Tris pH 7.2 at 1 mL/min. Remaining monomer concentration was measured from the area under the peak and calculated using a standard curve of known concentrations. The average remaining monomer is numerically shown. (D.) Lag time (T_lag_) and (E.) time to reach 50 % of maximum aggregation (t_50_) could only be calculated for assays containing calcium as there was no clear aggregation plateau in aSyn samples without calcium. Measurements were repeated with at least four sample replicates of three experiments and a one-way ANOVA with Sidak’s multiple comparison was used to calculate statistical significance. Error bars represent SEM.

**Figure S5.**
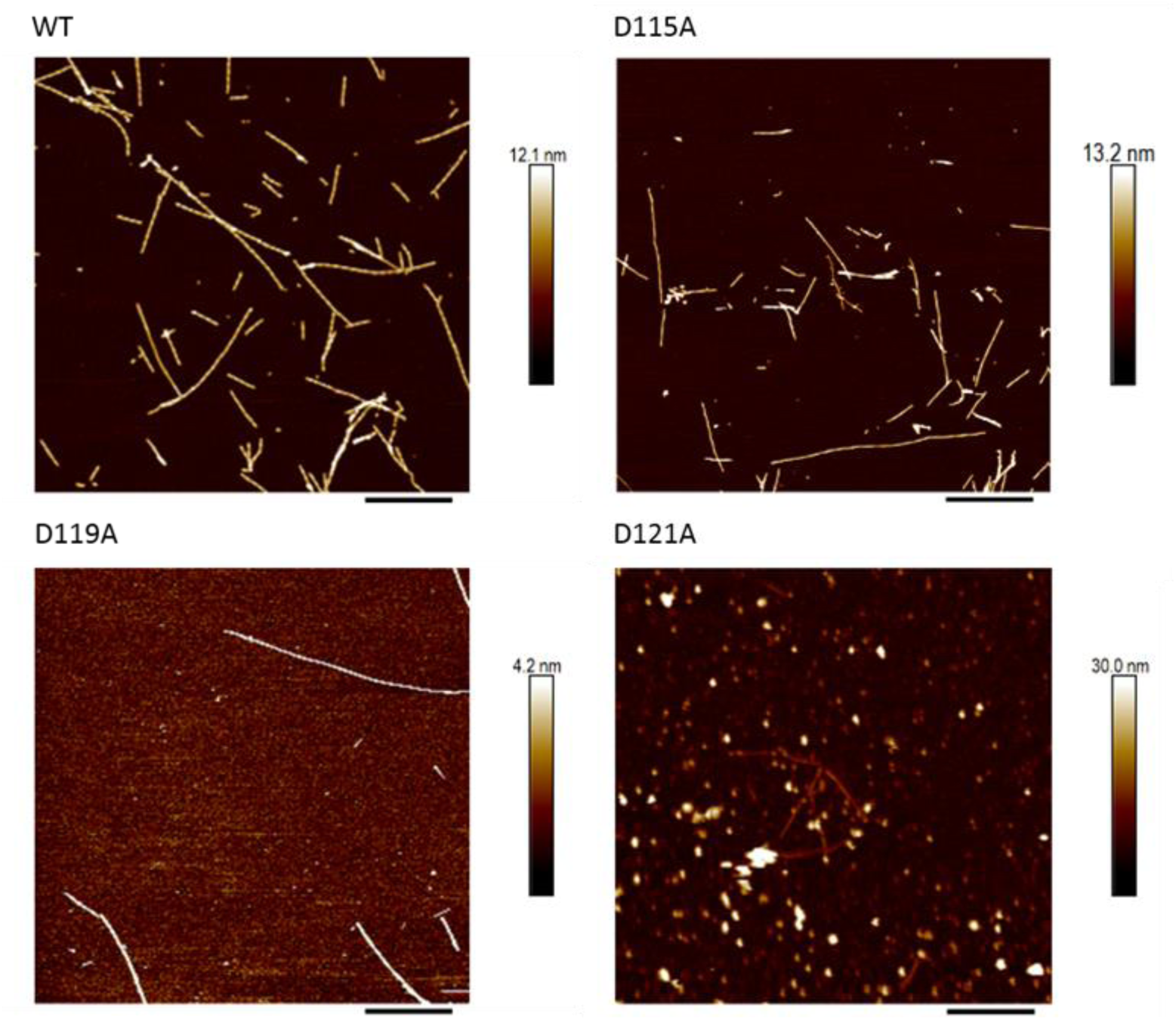
AFM reveals D121A aSyn forms mostly oligomeric structures, while WT, D115A and D119A aSyn form fibrillary structures upon incubation. aSyn samples were taken after ThT-based assays and incubated on freshly cleaved mica coated in 0.01% poly-lysine. Samples were washed in dH_2_O and dried. Representative images show fibril formation for WT, D115A and D119A aSyn samples, but mostly oligomeric structures formed in D121A aSyn samples. Scale bar = 800 nm.

**Figure S6.**
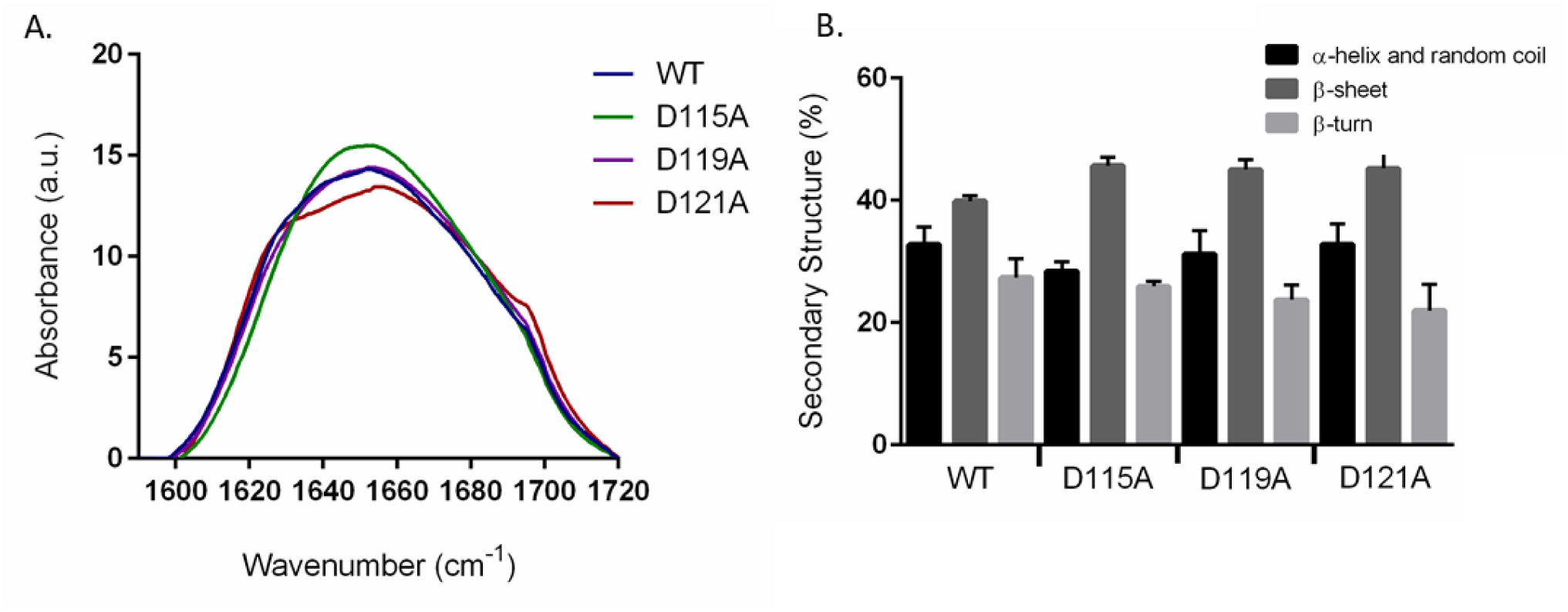
FTIR spectra and secondary structure analysis of WT and D to A mutant aSyn reveal no structural differences between the variants. 69 μM aSyn in 20 uM ammonium acetate was lyophilised and analysed using FTIR. (A) FTIR spectra of the absorbance of the aSyn D to A mutants, D115A (green), D119A (purple), D121A (red) and WT aSyn (blue) between 1590 - 1710 cm^-1^ and (B) their respective secondary structure as % distribution after spectra deconvolution. Three scans were performed for each sample and the experiment was repeated twice. A one-way ANOVA with Sidak’s multiple comparison was used to calculate statistical significance. Error bars represent SEM.

**Figure S7.**
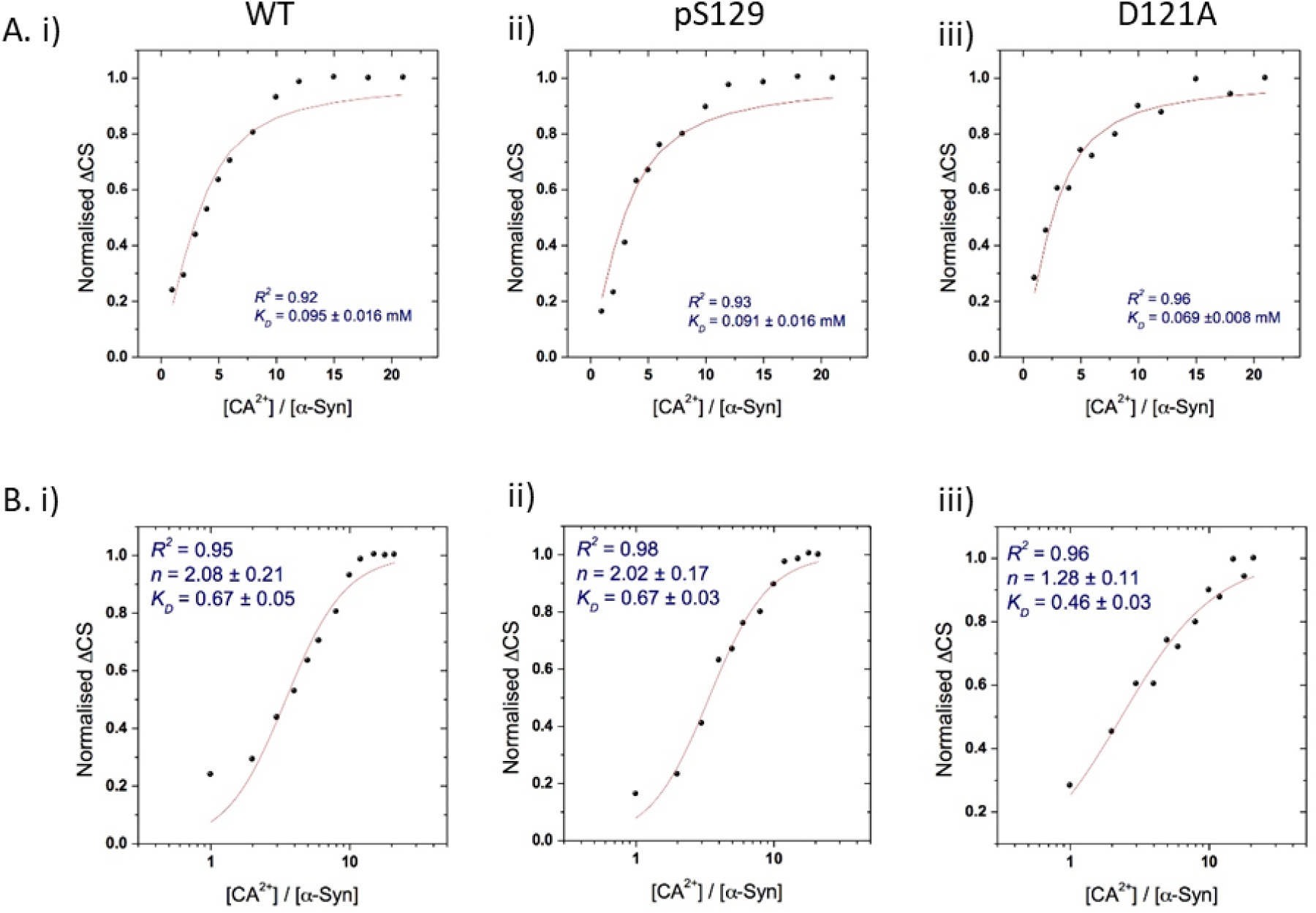
Dissociation constants of WT, pS129 and D121A aSyn and calcium reveal no significant differences between the different variants compared to WT aSyn and cooperative binding of calcium to aSyn. ^1^H-^15^N HSQC spectra of WT, pS129 and D121A aSyn (200 μM) was collected in the presence of increasing concentrations of Ca^2+^ (0.0 mM to 4.2 mM). (A) Fitting of aSyn calcium binding (*K_D_*), where the number of calcium ions bound is fixed to 3, based on MS data. Calculated *K_D_* were fitted for i) WT, ii) pS129, iii) D121A aSyn. (B) A second fitting was then used which takes into account cooperative binding using the Hill equation and gave *K_D_* values for i) WT, ii) pS129, iii) D121A aSyn. For all fittings using the Hill equation the *n* values are larger than 1 which shows positive cooperativity for the binding of calcium ions.

**Figure S8.**
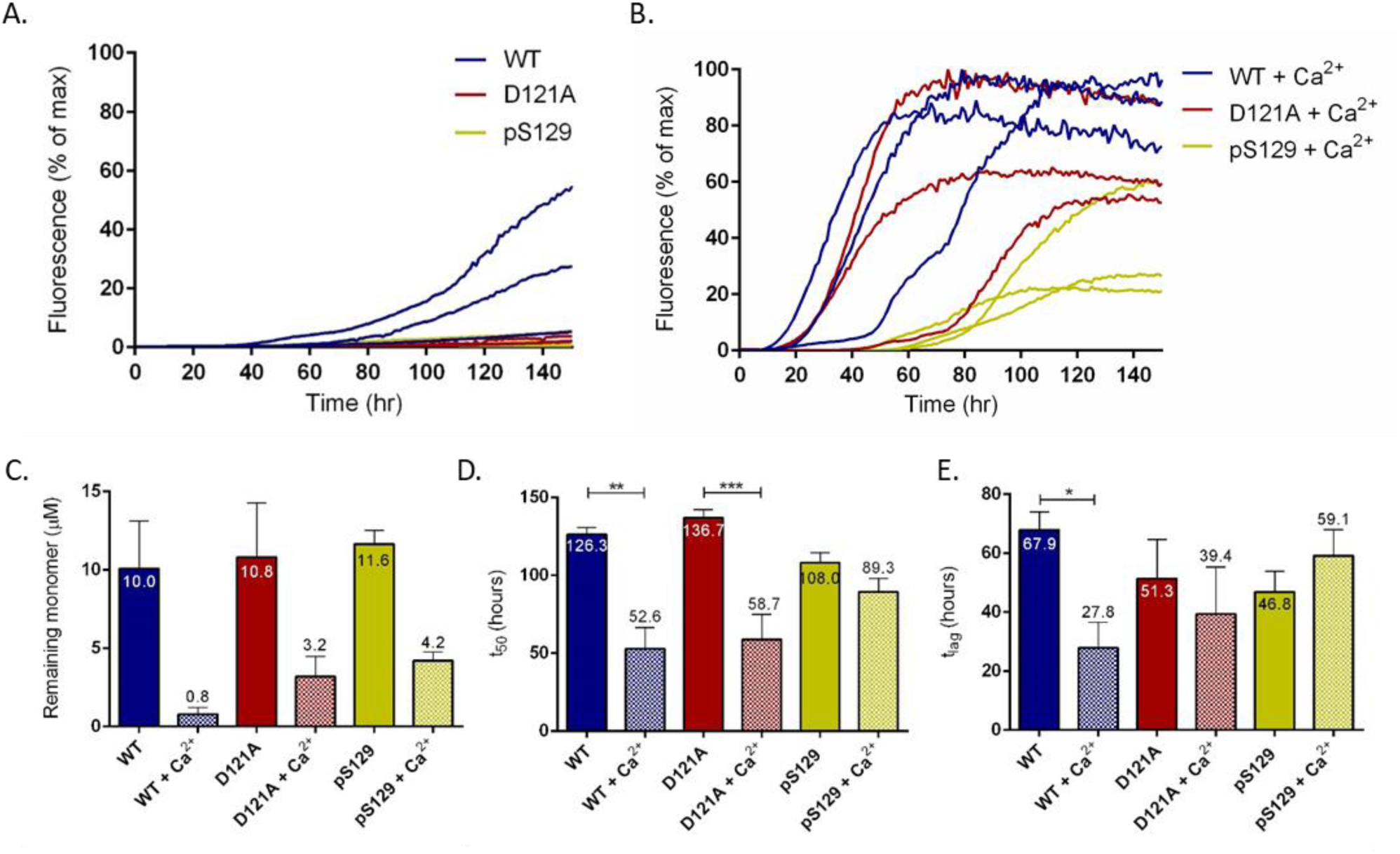
ThT-based aggregation assay reveals that D121A aSyn and pS129 aSyn do not aggregate as fast as WT aSyn. (A) The average of individual experiments are shown as ThT fluorescence intensity plotted as % of maximum fluorescence per plate of WT (blue), D121A (red) and pS129 aSyn (yellow) and (B) in the presence of 2.5 mM CaCl_2_. 20 μM aSyn was incubated with 20 μM in a half area 96 well plate with orbital agitation at 300 rpm for 5 minutes before each read every hour for 150 hours. (C.) Remaining monomer was measured using SEC-HPLC, 35 μL of monomer from each well in the ThT assay were analysed on an AdvanceBio SEC 130Å column equilibrated in 20 mM Tris pH 7.2 at 1 mL/min. Remaining monomer concentration was measured from the area under the peak and calculated using a standard curve of known concentrations. The average remaining monomer is numerically shown. (D.) Lag time (T_lag_) and (E.) time to reach 50 % of maximum aggregation (t_50_) were calculated and the average numerically shown. Measurements were repeated with at least four sample replicates of three experiments and a one-way ANOVA with Sidak’s multiple comparison was used to calculate statistical significance. Error bars represent SEM.

**Figure S9.**
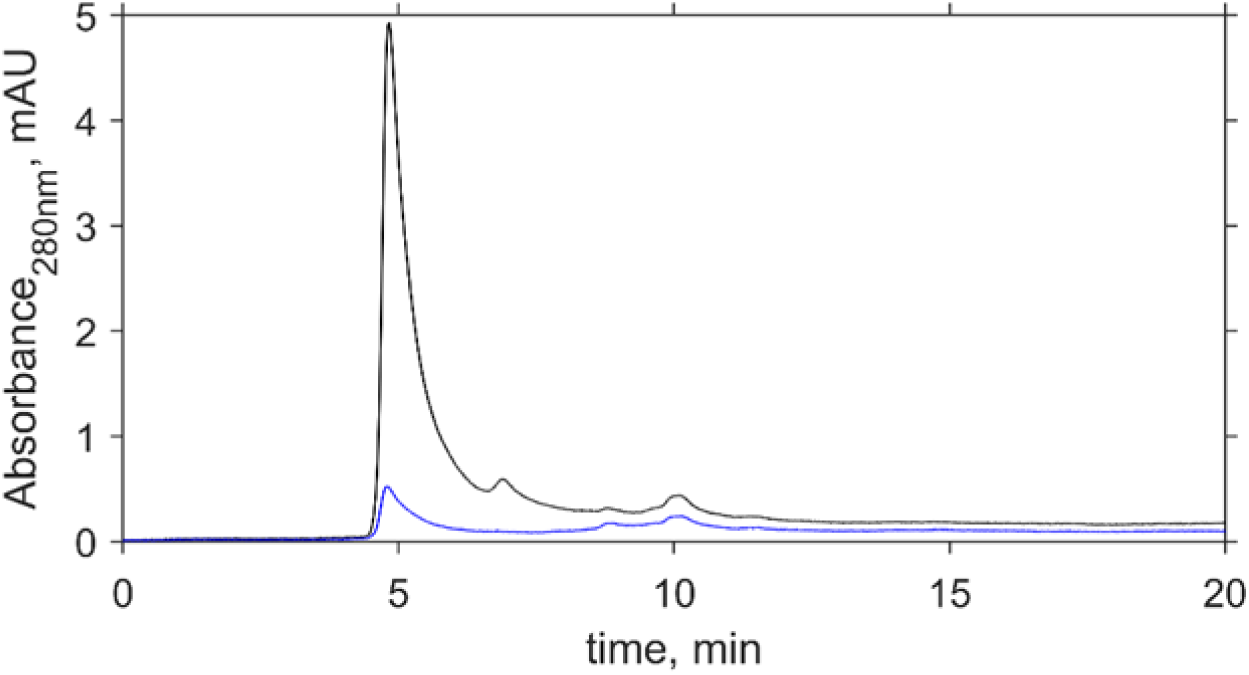
Representative chromatograph of the remaining aSyn monomer concentration as determined at the end of the ThT-based assay and analysed by size exclusion chromatography. aSyn samples taken from the wells of ThT-based assays were centrifuged to remove fibrils and 35 μL of the remaining monomer was analysed by HPLC-SEC. 35 μL was injected on to an AdvanceBio SEC 130Å column at a flow rate of 1 ml/min in 20 mM Tris pH 7.2 and absorbance measured at 280 nm. The area under the curve reflects the remaining monomer concentration, which is calculated using known protein standards. Representative remaining monomer concentration from one well of WT aSyn (black) and WT aSyn + 2.5 mM CaCl_2_ (blue) is shown.

**Figure S10.**
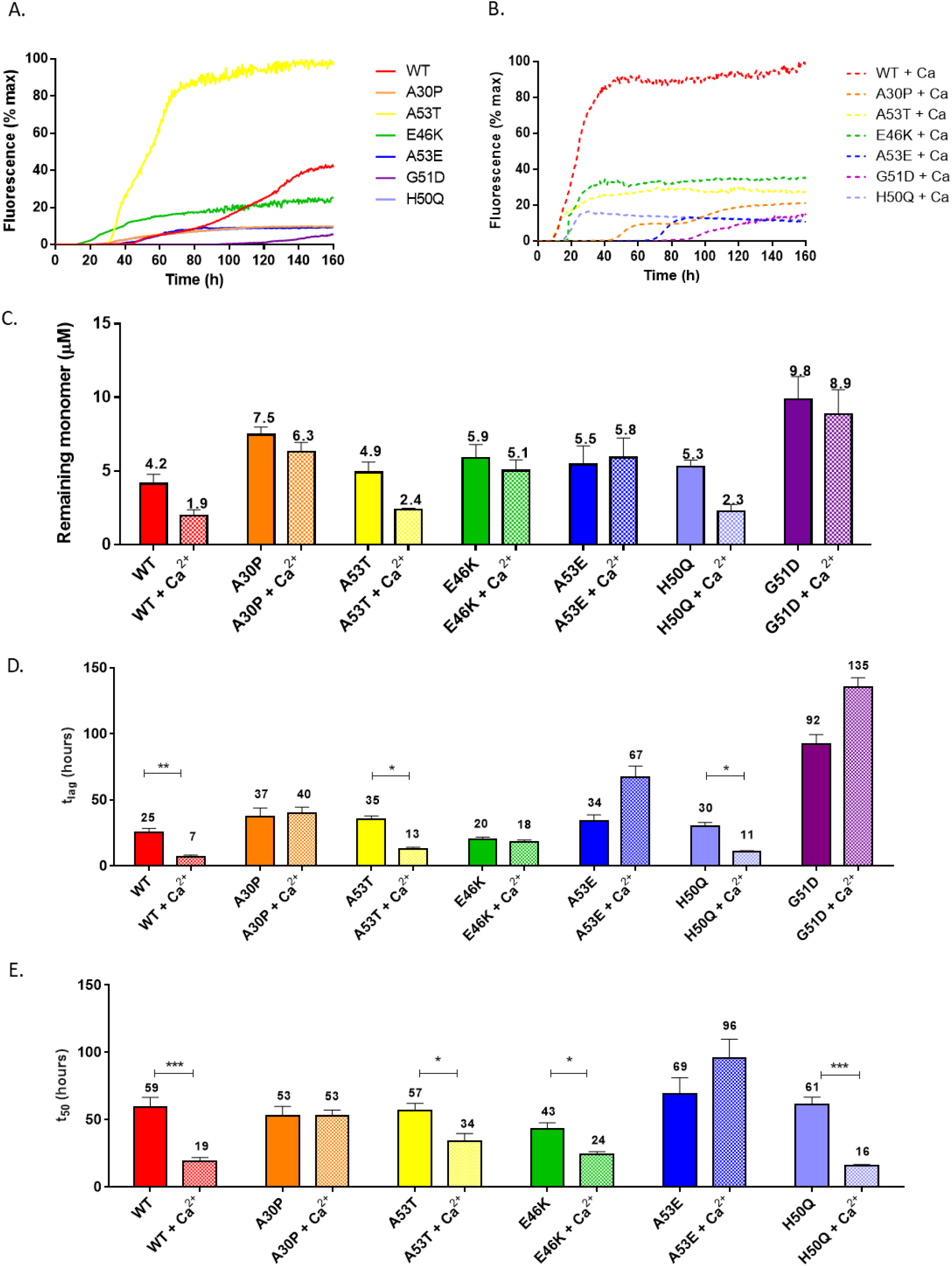
ThT-based aggregation assay reveals that the aSyn familial mutants display different aggregation behaviour upon the addition of calcium. (A) The average of individual experiments are shown as ThT fluorescence intensity plotted as % of maximum fluorescence per plate of WT (red), A30P (orange), A53T (yellow), E46K (green), A53E (blue), H50Q (lavender), G51D aSyn (purple) and (B) in the presence of 2.5 mM CaCl_2_. 20 μM aSyn was incubated with 20 μM in a half area 96 well plate with orbital agitation at 300 rpm for 5 minutes before each read every hour for 150 hours. (C.) Remaining monomer was measured using SEC-HPLC, 35 μL of monomer from each well in the ThT assays were analysed on an AdvanceBio SEC 130Å column equilibrated in 20 mM Tris pH 7.2 at 1 mL/min. Remaining monomer concentration was measured from the area under the peak and calculated using a standard curve of known concentrations. The average remaining monomer is numerically shown. (D.) Lag time (T_lag_) and (E.) time to reach 50 % of maximum aggregation (t_50_) were calculated and the average numerically shown. Mutant G51D aSyn did not reach the elongation phase in the studied timeframe, so the t_50_ value was not calculated. Measurements were repeated with at least four sample replicates of three experiments and a one-way ANOVA with Sidak’s multiple comparison was used to calculate statistical significance. Error bars represent SEM.

**Figure S11.**
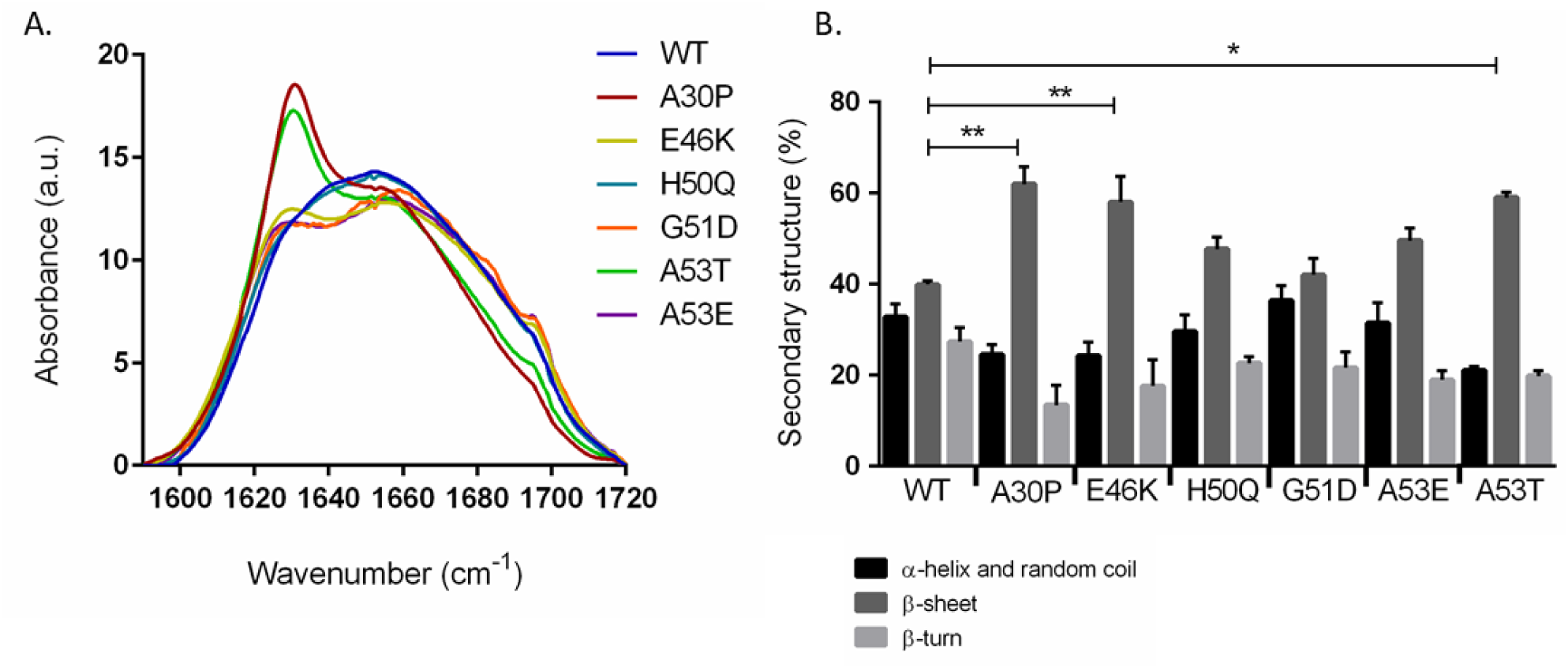
FTIR spectra and secondary structure analysis of monomeric WT aSyn and aSyn familial mutants reveal structural differences betweenthe aSyn variants. 69 μM aSyn in 20 μM ammonium acetate was lyophilised and analysed using FTIR. (A) FTIR spectra of the absorbance of WT aSyn (blue) and the aSyn familial mutants, A30P (red), E46K (yellow), H50Q (teal), G51D (orange), A53T (green), A53E aSyn (purple) between 1590 -1710 cm^-1^ and (B) their respective secondary structure as % distribution after spectra deconvolution. Three scans were performed for each sample and the experiments were repeated twice. A one-way ANOVA with Sidak’s multiple comparison was used to calculate statistical significance, for which p<0.05 * and p<0.01**. Error bars represent SEM. The % distribution of conformations was quantified by calculating the area under the curve for each peak in the chromatograph based on drift time (Figure 4b, Table S1, S2). It is likely that many conformations are present in each designated A-E conformation region as observed by the shoulders present in each peak as different conformations are not resolved (Figure S12)

**Figure S12.**
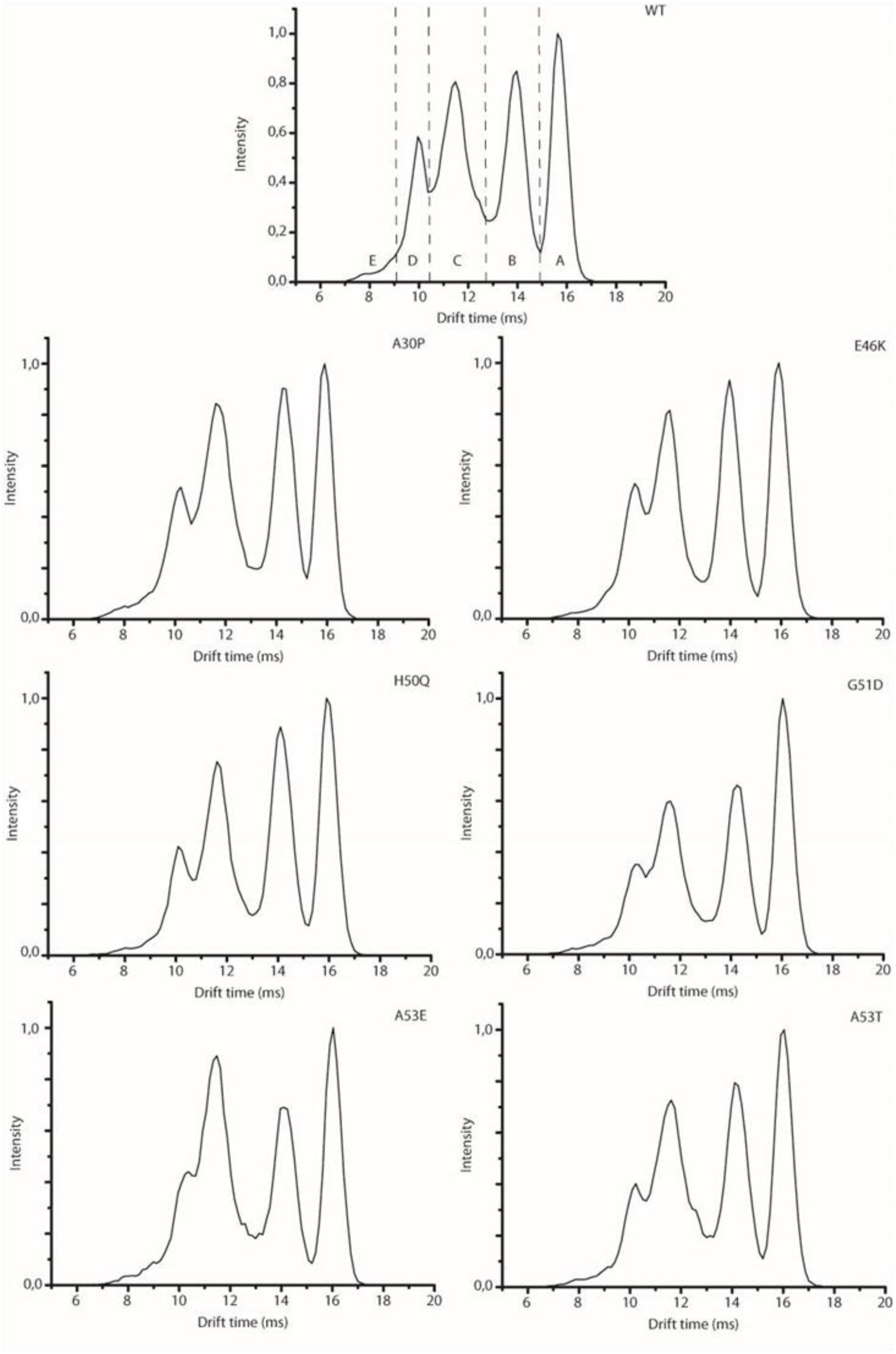
Arrival time distribution (ATD) of aSyn conformations. The average percentage distribution of the conformations was determined by identifying peaks in the ATD. Each peak was designated a letter A-E dependent on ATD (represented for WT aSyn). ATD is shown for WT, A30P, E46K, H50Q, G51D, A53E and A53T.

**Figure S13.**
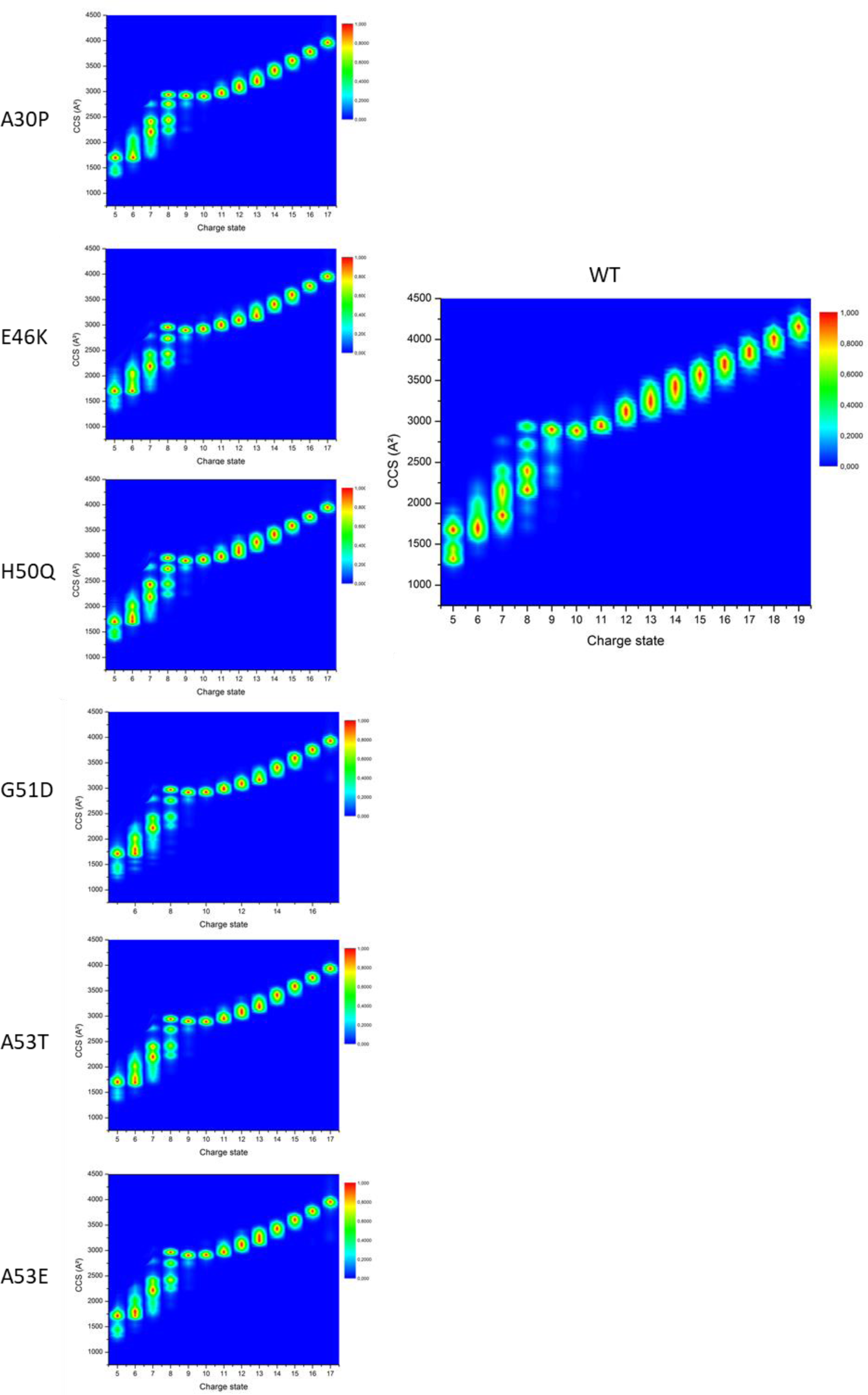
CCS values of aSyn WT and familial mutants show multiple coexisting conformations and extensions at higher charge states. Nano-ESI-IM-MS spectra of WT aSyn and aSyn familial mutants at all charge states detected (20 μM of aSyn in 50 mM ammonium acetate). Multiple defined conformations can be observed at charge states +9 and below. At the +10 charge state and above, charge repulsion leads to an extended, chain­like conformation as shown by the increasing CCS values.

**Table S1.** Average percentage of the distribution of conformations of WT aSyn and mutants determined by nano-ESI-IM-MS in the absence calcium shows small changes in percentage distribution of conformations between the aSyn and its variants.

| Conformation<br>without calcium | WT<br>av. % | A30P<br>av. % | E46K<br>av. % | A53E<br>av. % | A53T<br>av. % | H50Q<br>av. % | G51D<br>av. % |
| --- | --- | --- | --- | --- | --- | --- | --- |
| A | 23.54 | 21.86 | 26.22* | 27.29 | 26.57* | 25.96 | 28.05 |
| B | 26.80 | 26.48 | 27.72 | 24.97 | 26.78 | 30.47 | 24.92 |
| C | 33.09 | 35.74* | 29.46** | 34.38 | 32.88 | 30.26 | 30.49* |
| D | 15.58 | 14.83 | 15.74 | 12.12** | 12.62** | 12.37* | 14.38 |
| E | 0.98 | 1.10 | 0.85 | 1.20 | 1.10 | 0.92 | 2.14 |

**Table S2.** Average percentage of the distribution of conformations of WT and aSyn mutants determined by nano-ESI-IM-MS in the presence of calcium shows significant differences upon the addition of calcium to aSyn and its variants.

| Conformation<br>with calcium | WT<br>av. % | A30P<br>av. % | E46K<br>av. % | A53E<br>av. % | A53T<br>av. % | H50Q<br>av. % | G51D<br>av. % |
| --- | --- | --- | --- | --- | --- | --- | --- |
| A | 21.53 | 22.79 | 25.82* | 24.18 | 25.40* | 24.86 | 25.00 |
| B | 14.73 | 19.96** | 17.54** | 15.39 | 18.66** | 17.74** | 16.06 |
| C | 37.85 | 37.57 | 32.74* | 39.04 | 38.66 | 37.74 | 36.16 |
| D | 22.83 | 17.57* | 21.58 | 18.93* | 15.96** | 18.06 | 20.10 |
| E | 3.07 | 2.10* | 2.32 | 2.45 | 1.84* | 2.16 | 2.68 |

\*p<0.05
\*\*p<0.01

**Figure S14.**
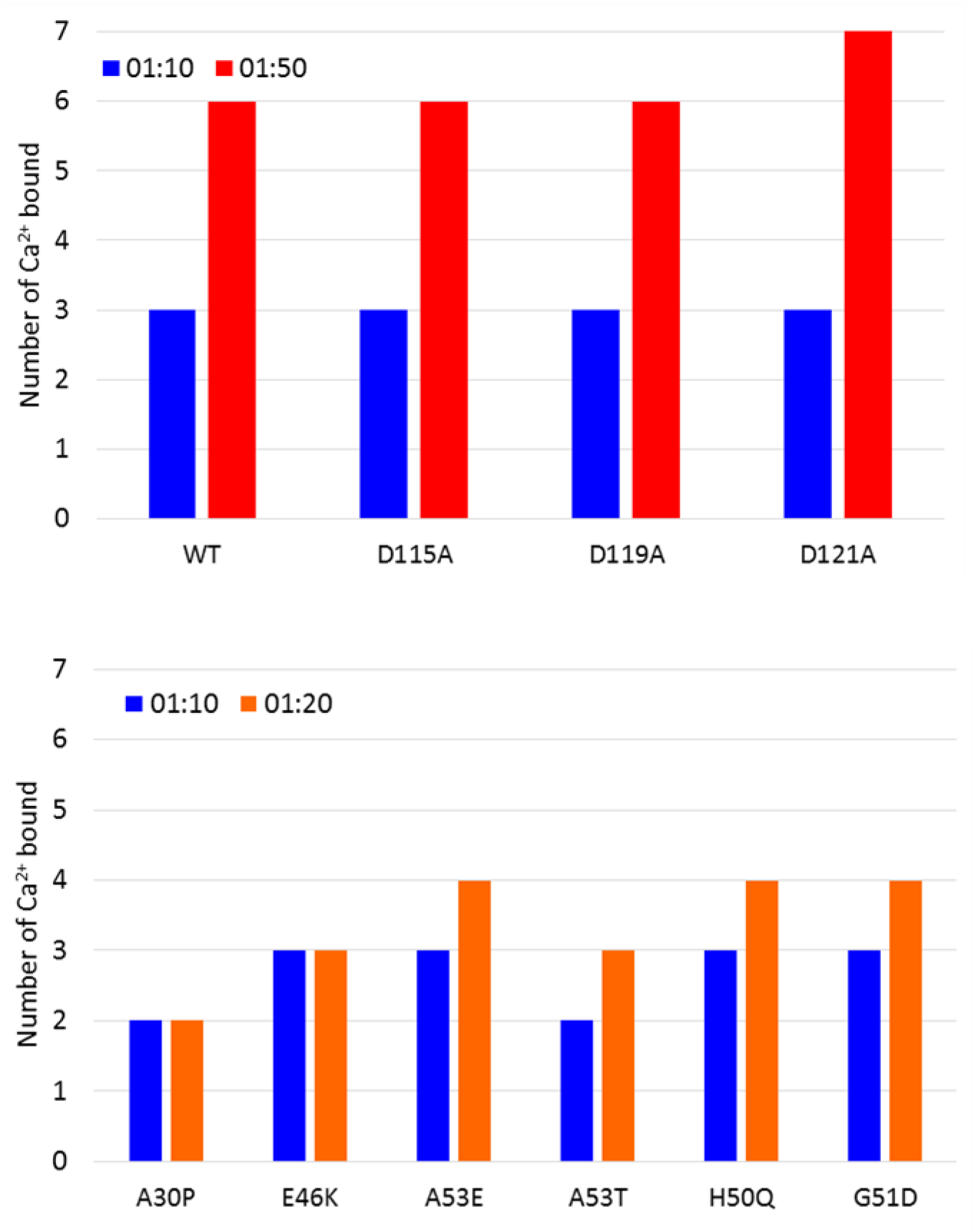
Analysis of number of Ca^2+^ ions bound to different aSyn mutants at different protein to ratios reveals no significant differences between the different aSyn variants. The number of calcium ions bound to 20 μM of aSyn in 20 mM ammonium acetate at a protein to calcium ratio of 1:10 (blue) 1:20 (orange) and 1:50 (red) was determined from the mass of the different metallated species in the native nano ESI-MS spectra.

**Figure S15.**
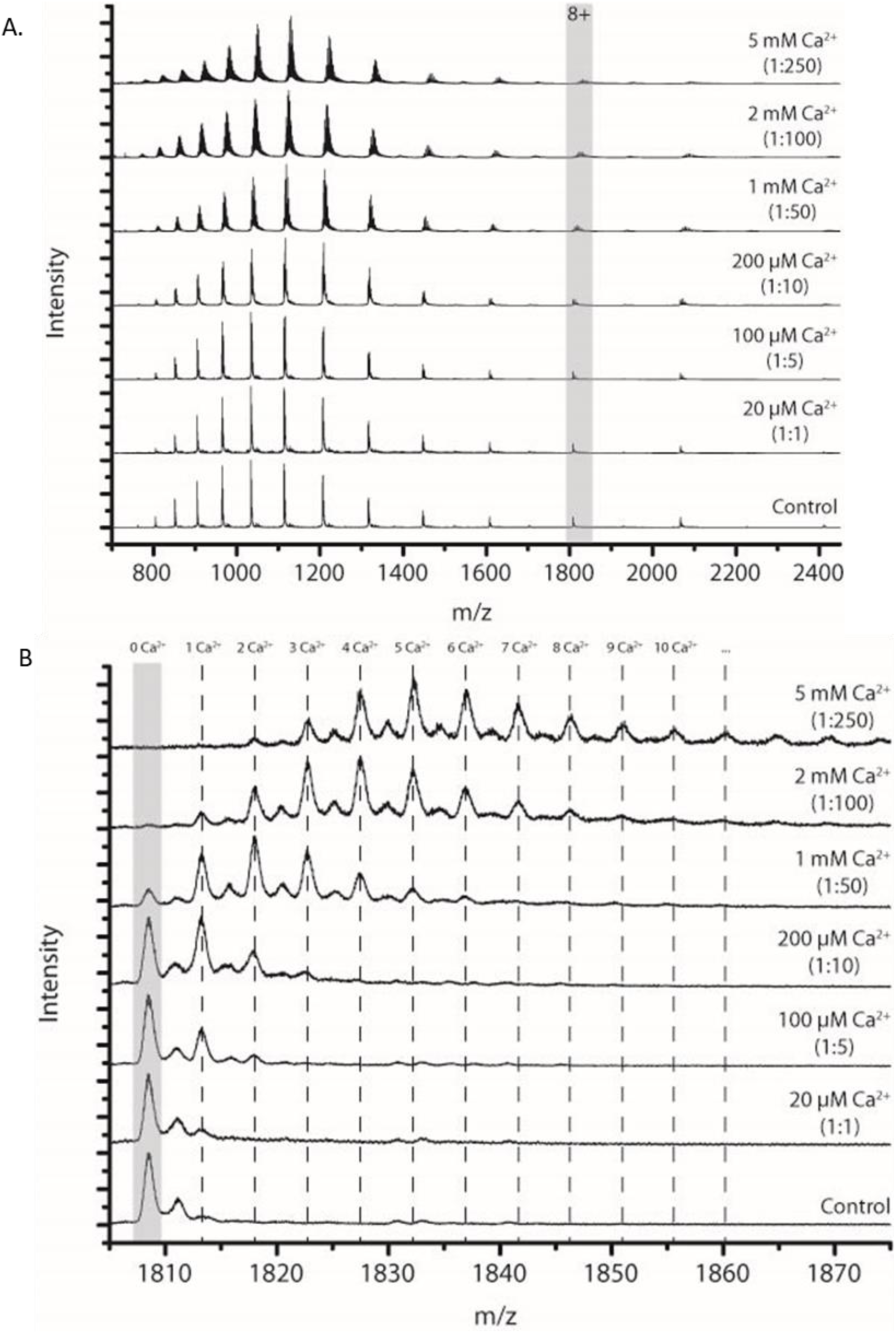
Native nano ESI-MS spectra of WT aSyn with increasing concentrations of calcium shows that up to 11 calcium ions can be bound to aSyn at a 1:250 protein to calcium ratio. (A) The native nano ESI-MS full spectrum of WT aSyn with calcium in ratios 1:1,1:5,1:10,1:50, 1:100 and 1:250. (B) The zoomed in 8+ charge state of aSyn shows that at least 11 calcium ions can be bound to aSyn in a 1:250 protein to calcium ratio. 20 μM aSyn in 20 mM ammonium acetate (pH 7) was used in all experiments.

## Methods and Materials for Supplementary Data

### Thioflavin-T (ThT) based assays

10 μM freshly made ThT (abeam, Cambridge, UK) was added to 50 μL of 10 μM aSyn in 20 mM Tris pH 7.2. All samples were loaded onto nonbinding, clear bottom, 96-well half-area plates (Greiner Bio-One GmbH, Germany). The plates were sealed with a SILVERseal aluminium microplate sealer (Grenier Bio-One GmbH). Fluorescence measurements were taken with a FLUOstar Omega plate reader (BMG LABTECH GmbH, Ortenbery, Germany). The plates were incubated at 37°C with orbital shaking at 300 rpm for five minutes before each read every hour. Excitation was set at 440 nm with 20 flashes and the ThT fluorescence intensity measured at 480 nm emission with a 1300 gain setting. ThT assays were repeated at least 3 times using four or more wells for each condition. Data were normalised to the well with the maximum fluorescence intensity for each plate and the average fluorescence intensity was calculated for all experiments. The lag time (tlag) and half life of the fluorescence (t50) were calculated for each aSyn variant.

### Analytical Size Exclusion Chromatography (SEC) of remaining monomer after ThT assays

SEC-HPLC analysis was used to calculate the remaining aSyn monomer concentration in each well at the end of the ThT assays. The contents of each well after the ThT-based assay were centrifuged at 21k x g for 20 minutes and the supernatant added to individual aliquots in the autosampler of the Agilent 1260 Infinity HPLC system (Agilent Technologies LDA UK Limited, UK). 35 pL of each sample was injected onto an Advance Bio SEC column, 7.8 × 300 mm 300Å (Agilent, UK) in 20 mM Tris pH 7.2 at 1 mL/min flow-rate. The elution profile was monitored by UV absorption at 220 and 280 nm. A calibration curve of known concentrations of aSyn was used to calculate the remaining monomer concentration of aSyn in each well. Two or three wells per experiment for three experiments were analysed for the remaining monomer concentration.

### Fourier transform infrared spectroscopy (FTIR)

To prepare samples for FTIR analysis, aSyn monomer was buffer exchanged into 20 mM ammonium acetate pH 7 using PD10 Desalting columns (GE Healthcare). The samples were snap frozen in liquid nitrogen and lyophilised in a LyoQuest 85 freeze-dryer (Telstar, Spain). ∼300 pg protein was mixed with ∼100 mg KBr using an agate mortar and pressed to 10 tonnes to form self-supporting disks. The FTIR spectra were collected on a Cary 680 FTIR spectrometer with 60 scans and a resolution of 1 cm^-1^. The FTIR spectra were plotted by subtracting the KBr background spectrum, performing a spline baseline correction, and normalising the spectra to the area under the curve.

### Atomic Force Microscopy (AFM)

The contents of wells from the ThT-based assays were centrifuged for 20 minutes at 21 k x g. The supernatant was removed to keep 10 pL with remaining fibrils. The fibrils were resuspended and incubated on a freshly cleaved mica surface, which had been coated in 0.1% poly-l-lysine, for 20 min. The mica was washed three times in 18.2 Ω dH_2_O to remove lose protein. Images were acquired in dH_2_O using tapping mode on a BioScope Resolve (Bruker, AXS GmBH) using ScanAsyst-Fluid+ probes. 512 lines were acquired at a scan rate of 1.5-2 Hz per image with a field of view of 2-5 μm and for at least six fields of view. Images were adjusted for contrast and exported from NanoScope Analysis 8.2 software (Bruker).

## References

(1) Shahmoradian, S. H.; Lewis, A. J.; Genoud, C.; Hench, J.; Moors, T. E.; Navarro, P. P.; Castaño-Díez, D.; Schweighauser, G.; Graff-Meyer, A.; Goldie, K. N.;, et al. Lewy Pathology in Parkinson’s Disease Consists of Crowded Organelles and Lipid Membranes. Nat. Neurosci. 2019, 22 (7), 1099–1109. https://doi.org/10.1038/s41593-019-0423-2.

(2) Mahul-Mellier, A.-L.; Altay, M. F.; Burtscher, J.; Maharjan, N.; Ait-Bouziad, N.; Chiki, A.; Vingill, S.; Wade-Martins, R.; Holton, J.; Strand, C.;, et al. The Making of a Lewy Body: The Role of α-Synuclein Post-Fibrillization Modifications in Regulating the Formation and the Maturation of Pathological Inclusions. bioRxiv 2018, 500058. https://doi.org/10.1101/500058.

(3) Lautenschläger, J.; Kaminski, C. F.; Kaminski Schierle, G. S. α-Synuclein - Regulator of Exocytosis, Endocytosis, or Both? Trends Cell Biol. 2017. https://doi.org/10.1016/j.tcb.2017.02.002.

(4) Logan, T.; Bendor, J.; Toupin, C.; Thorn, K.; Edwards, R. H. α-Synuclein Promotes Dilation of the Exocytotic Fusion Pore. Nat. Publ. Gr. 2017. https://doi.org/10.1038/nn.4529.

(5) Uversky, V. N.; Li, J.; Souillac, P.; Millett, I. S.; Doniach, S.; Jakes, R.; Goedert, M.; Fink, A. L. Biophysical Properties of the Synucleins and Their Propensities to Fibrillate. J. Biol. Chem. 2002, 277 (14), 11970–11978. https://doi.org/10.1074/jbc.M109541200.

(6) Stephens, A. D.; Zacharopoulou, M.; Kaminski Schierle, G. S. The Cellular Environment Affects Monomeric α-Synuclein Structure. Trends Biochem. Sci. 2019, 44 (5), 453–466. https://doi.org/10.1016/j.tibs.2018.11.005.

(7) Giasson, B. I.; Murray, I. V; Trojanowski, J. Q.; Lee, V. M. A Hydrophobic Stretch of 12 Amino Acid Residues in the Middle of Alpha-Synuclein Is Essential for Filament Assembly. J. Biol. Chem. 2001, 276 (4), 2380–2386. https://doi.org/10.1074/jbc.M008919200.

(8) Binolfi, A.; Rasia, R. M.; Bertoncini, C. W.; Ceolin, M.; Zweckstetter, M.; Griesinger, C.; Jovin, T. M.; Fernández, C. O. Interaction of α-Synuclein with Divalent Metal Ions Reveals Key Differences: A Link between Structure, Binding Specificity and Fibrillation Enhancement. J. Am. Chem. Soc. 2006, 128 (30), 9893–9901. https://doi.org/10.1021/ja0618649.

(9) Polymeropoulos, M. H.; Lavedan, C.; Leroy, E.; Ide, S. E.; Dehejia, A.; Dutra, A.; Pike, B.; Root, H.; Rubenstein, J.; Boyer, R.;, et al. Mutation in the α-Synuclein Gene Identified in Families with Parkinson’s Disease. Science (80-). 1997, 276 (5321), 2045–2047. https://doi.org/10.1126/science.276.5321.2045.

(10) Krüger, R.; Kuhn, W.; Müller, T.; Woitalla, D.; Graeber, M.; Kösel, S.; Przuntek, H.; Epplen, J. T.; Schols, L.; Riess, O. Ala30Pro Mutation in the Gene Encoding α-Synuclein in Parkinson’s Disease. Nat. Genet. 1998, 18 (2), 106–108. https://doi.org/10.1038/ng0298-106.

(11) Zarranz, J. J.; Alegre, J.; Gómez-Esteban, J. C.; Lezcano, E.; Ros, R.; Ampuero, I.; Vidal, L.; Hoenicka, J.; Rodriguez, O.; Atarés, B.;, et al. The New Mutation, E46K, of α-Synuclein Causes Parkinson and Lewy Body Dementia. Ann. Neurol. 2004, 55 (2), 164–173. https://doi.org/10.1002/ana.10795.

(12) Proukakis, C.; Dudzik, C. G.; Brier, T.; MacKay, D. S.; Cooper, J. M.; Millhauser, G. L.; Houlden, H.; Schapira, A. H. A Novel α-Synuclein Missense Mutation in Parkinson Disease. Neurology. American Academy of Neurology March 12, 2013, pp 1062–1064. https://doi.org/10.1212/WNL.0b013e31828727ba.

(13) Appel-Cresswell, S.; Vilarino-Guell, C.; Encarnacion, M.; Sherman, H.; Yu, I.; Shah, B.; Weir, D.; Thompson, C.; Szu-Tu, C.; Trinh, J.;, et al. Alpha-Synuclein p.H50Q, a Novel Pathogenic Mutation for Parkinson’s Disease. Mov. Disord. 2013, 28 (6), 811–813. https://doi.org/10.1002/mds.25421.

(14) Lesage, S.; Anheim, M.; Letournel, F.; Bousset, L.; Honoré, A.; Rozas, N.; Pieri, L.; Madiona, K.; Dürr, A.; Melki, R.;, et al. G51D α-Synuclein Mutation Causes a Novel Parkinsonian-Pyramidal Syndrome. Ann. Neurol. 2013, 73 (4), 459–471. https://doi.org/10.1002/ana.23894.

(15) Pasanen, P.; Myllykangas, L.; Siitonen, M.; Raunio, A.; Kaakkola, S.; Lyytinen, J.; Tienari, P. J.; Pöyhönen, M.; Paetau, A. A Novel α-Synuclein Mutation A53E Associated with Atypical Multiple System Atrophy and Parkinson’s Disease-Type Pathology. Neurobiol. Aging 2014, 35 (9), 2180.e1-2180.e5. https://doi.org/10.1016/j.neurobiolaging.2014.03.024.

(16) Lill, C. M. Genetics of Parkinson’s Disease. Mol. Cell. Probes 2016, 30 (6), 386–396. https://doi.org/10.1016/j.mcp.2016.11.001.

(17) Wu, K. P.; Baum, J. Detection of Transient Interchain Interactions in the Intrinsically Disordered Protein α-Synuclein by NMR Paramagnetic Relaxation Enhancement. J. Am. Chem. Soc. 2010, 132 (16), 5546–5547. https://doi.org/10.1021/ja9105495.

(18) Stephens, A.; Nespovitaya, N.; Zacharopoulou, M.; Kaminski, F.; Phillips, J. J.; Schierle, G. S. K. Different Structural Conformers of Monomeric Alpha-Synuclein Identified after Lyophilising and Freezing. Anal. Chem. 2018, acs.analchem.8b01264. https://doi.org/10.1021/acs.analchem.8b01264.

(19) Dedmon, M. M.; Lindorff-Larsen, K.; Christodoulou, J.; Vendruscolo, M.; Dobson, C. M. Mapping Long-Range Interactions in α-Synuclein Using Spin-Label NMR and Ensemble Molecular Dynamics Simulations. J. Am. Chem. Soc. 2005, 127 (2), 476–477. https://doi.org/10.1021/ja044834j.

(20) Zhou, W.; Long, C.; Reaney, S. H.; Di Monte, D. A.; Fink, A. L.; Uversky, V. N. Methionine Oxidation Stabilizes Non-Toxic Oligomers of α-Synuclein through Strengthening the Auto-Inhibitory Intra-Molecular Long-Range Interactions. Biochim. Biophys. Acta - Mol. Basis Dis. 2010, 1802 (3), 322–330. https://doi.org/10.1016/j.bbadis.2009.12.004.

(21) Esteban-Martín, S.; Silvestre-Ryan, J.; Bertoncini, C. W.; Salvatella, X. Identification of Fibril-like Tertiary Contacts in Soluble Monomeric α-Synuclein. Biophys. J. 2013, 105 (5), 1192– 1198. https://doi.org/10.1016/j.bpj.2013.07.044.

(22) McClendon, S.; Rospigliosi, C. C.; Eliezer, D. Charge Neutralization and Collapse of the C-Terminal Tail of Alpha-Synuclein at Low PH. Protein Sci. 2009, 18 (7), 1531–1540. https://doi.org/10.1002/pro.149.

(23) Ranjan, P.; Kumar, A. Perturbation in Long-Range Contacts Modulates the Kinetics of Amyloid Formation in α-Synuclein Familial Mutants. ACS Chem. Neurosci. 2017, 8 (10), 2235–2246. https://doi.org/10.1021/acschemneuro.7b00149.

(24) Sung, Y.; Eliezer, D. Residual Structure, Backbone Dynamics, and Interactions within the Synuclein Family. J. Mol. Biol. 2007, 372 (3), 689–707. https://doi.org/10.1016/J.JMB.2007.07.008.

(25) Bertoncini, C. W.; Jung, Y.-S.; Fernandez, C. O.; Hoyer, W.; Griesinger, C.; Jovin, T. M.; Zweckstetter, M. From The Cover: Release of Long-Range Tertiary Interactions Potentiates Aggregation of Natively Unstructured -Synuclein. Proc. Natl. Acad. Sci. 2005, 102 (5), 1430– 1435. https://doi.org/10.1073/pnas.0407146102.

(26) Afitska, K.; Fucikova, A.; Shvadchak, V. V; Yushchenko, D. A. Modification of C Terminus Provides New Insights into the Mechanism of α-Synuclein Aggregation. Biophys. J. 2017, 113 (10), 2182–2191. https://doi.org/10.1016/j.bpj.2017.08.027.

(27) Ulrih, N. P.; Barry, C. H.; Fink, A. L. Impact of Tyr to Ala Mutations on α-Synuclein Fibrillation and Structural Properties. Biochim. Biophys. Acta - Mol. Basis Dis. 2008, 1782 (10), 581–585. https://doi.org/10.1016/j.bbadis.2008.07.004.

(28) Landeck, N.; Strathearn, K. E.; Ysselstein, D.; Buck, K.; Hulleman, J. D.; Hindupur, J.; Griggs, A. M.; Padalkar, S.; Stanciu, A.; Kirik, D.;, et al. Two C-Terminal Sequence Variations Determine Differential Neurotoxicity between Human and Mouse α -Synuclein Natalie Landeck, 1, Katherine E. Strathearn, 2,. bioRxiv 2019, 700377. https://doi.org/10.1101/700377.

(29) Roeters, S. J.; Iyer, A.; Pletikapić, G.; Kogan, V.; Subramaniam, V.; Woutersen, S. Evidence for Intramolecular Antiparallel Beta-Sheet Structure in Alpha-Synuclein Fibrils from a Combination of Two-Dimensional Infrared Spectroscopy and Atomic Force Microscopy. Sci. Rep. 2017, 7, 41051. https://doi.org/10.1038/srep41051.

(30) Anderson, J. P.; Walker, D. E.; Goldstein, J. M.; de Laat, R.; Banducci, K.; Caccavello, R. J.; Barbour, R.; Huang, J.; Kling, K.; Lee, M.;, et al. Phosphorylation of Ser-129 Is the Dominant Pathological Modification of Alpha-Synuclein in Familial and Sporadic Lewy Body Disease. J. Biol. Chem. 2006, 281 (40), 29739–29752. https://doi.org/10.1074/jbc.M600933200.

(31) Fujiwara, H.; Hasegawa, M.; Dohmae, N.; Kawashima, A.; Masliah, E.; Goldberg, M. S.; Shen, J.; Takio, K.; Iwatsubo, T. Α-Synuclein Is Phosphorylated in Synucleinopathy Lesions. Nat. Cell Biol. 2002, 4 (2), 160–164. https://doi.org/10.1038/ncb748.

(32) Samuel, F.; Flavin, W. P.; Iqbal, S.; Pacelli, C.; Renganathan, S. D. S.; Trudeau, L. E.; Campbell, E. M.; Fraser, P. E.; Tandon, A. Effects of Serine 129 Phosphorylation on α-Synuclein Aggregation, Membrane Association, and Internalization. J. Biol. Chem. 2016, 291 (9), 4374– 4385. https://doi.org/10.1074/jbc.M115.705095.

(33) Paleologou, K. E.; Schmid, A. W.; Rospigliosi, C. C.; Kim, H. Y.; Lamberto, G. R.; Fredenburg, R. A.; Lansbury, P. T.; Fernandez, C. O.; Eliezer, D.; Zweckstetter, M.;, et al. Phosphorylation at Ser-129 but Not the Phosphomimics S129E/D Inhibits the Fibrillation of α-Synuclein. J. Biol. Chem. 2008, 283 (24), 16895–16905. https://doi.org/10.1074/jbc.M800747200.

(34) Bertoncini, C. W.; Fernandez, C. O.; Griesinger, C.; Jovin, T. M.; Zweckstetter, M. Familial Mutants of α-Synuclein with Increased Neurotoxicity Have a Destabilized Conformation. J. Biol. Chem. 2005, 280 (35), 30649–30652. https://doi.org/10.1074/jbc.C500288200.

(35) Bhattacharyya, D.; Kumar, R.; Mehra, S.; Ghosh, A.; Maji, S. K.; Bhunia, A. Multitude NMR Studies of α-Synuclein Familial Mutants: Probing Their Differential Aggregation Propensities. Chem. Commun. 2018, 54 (29), 3605–3608. https://doi.org/10.1039/c7cc09597j.

(36) Flagmeier, P.; Meisl, G.; Vendruscolo, M.; Knowles, T. P. J.; Dobson, C. M.; Buell, A. K.; Galvagnion, C. Mutations Associated with Familial Parkinson’s Disease Alter the Initiation and Amplification Steps of α-Synuclein Aggregation. Proc. Natl. Acad. Sci. 2016, 113 (37), 10328–10333. https://doi.org/10.1073/pnas.1604645113.

(37) Wise-Scira, O.; Dunn, A.; Aloglu, A. K.; Sakallioglu, I. T.; Coskuner, O. Structures of the E46K Mutant-Type α-Synuclein Protein and Impact of E46K Mutation on the Structures of the Wild-Type α-Synuclein Protein. ACS Chem. Neurosci. 2013, 4 (3), 498–508. https://doi.org/10.1021/cn3002027.

(38) Santner, A.; Uversky, V. N. Metalloproteomics and Metal Toxicology of α-Synuclein. Metallomics 2010, 2 (6), 378–392. https://doi.org/10.1039/b926659c.

(39) Lautenschläger, J.; Stephens, A. D.; Fusco, G.; Ströhl, F.; Curry, N.; Zacharopoulou, M.; Michel, C. H.; Laine, R.; Nespovitaya, N.; Fantham, M.;, et al. C-Terminal Calcium Binding of α-Synuclein Modulates Synaptic Vesicle Interaction. Nat. Commun. 2018, 9 (1), 712. https://doi.org/10.1038/s41467-018-03111-4.

(40) Surmeier, D. J.; Guzman, J. N.; Sanchez-Padilla, J. Calcium, Cellular Aging, and Selective Neuronal Vulnerability in Parkinson’s Disease. Cell Calcium 2010, 47 (2), 175–182. https://doi.org/10.1016/J.CECA.2009.12.003.

(41) Lu, Y.; Prudent, M.; Fauvet, B.; Lashuel, H. A.; Girault, H. H. Phosphorylation of α-Synuclein at Y125 and S129 Alters Its Metal Binding Properties: Implications for Understanding the Role of α-Synuclein in the Pathogenesis of Parkinson’s Disease and Related Disorders. ACS Chem. Neurosci. 2011, 2 (11), 667–675. https://doi.org/10.1021/cn200074d.

(42) Ghosh, D.; Sahay, S.; Ranjan, P.; Salot, S.; Mohite, G. M.; Singh, P. K.; Dwivedi, S.; Carvalho, E.; Banerjee, R.; Kumar, A.;, et al. The Newly Discovered Parkinsons Disease Associated Finnish Mutation (A53E) Attenuates α-Synuclein Aggregation and Membrane Binding. Biochemistry 2014, 53 (41), 6419–6421. https://doi.org/10.1021/bi5010365.

(43) Galvagnion, C.; Buell, A. K.; Meisl, G.; Michaels, T. C. T.; Vendruscolo, M.; Knowles, T. P. J.; Dobson, C. M. Lipid Vesicles Trigger α-Synuclein Aggregation by Stimulating Primary Nucleation. Nat. Chem. Biol. 2015, 11 (3), 229–234. https://doi.org/10.1038/nchembio.1750.

(44) Wongkongkathep, P.; Han, J. Y.; Choi, T. S.; Yin, S.; Kim, H. I.; Loo, J. A. Native Top-Down Mass Spectrometry and Ion Mobility MS for Characterizing the Cobalt and Manganese Metal Binding of α-Synuclein Protein. J. Am. Soc. Mass Spectrom. 2018, 29 (9), 1870–1880. https://doi.org/10.1007/s13361-018-2002-2.

(45) Konijnenberg, A.; Ranica, S.; Narkiewicz, J.; Legname, G.; Grandori, R.; Sobott, F.; Natalello, A. Opposite Structural Effects of Epigallocatechin-3-Gallate and Dopamine Binding to α-Synuclein. Anal. Chem. 2016, 88 (17), 8468–8475. https://doi.org/10.1021/acs.analchem.6b00731.

(46) Brodie, N. I.; Popov, K. I.; Petrotchenko, E. V.; Dokholyan, N. V.; Borchers, C. H. Conformational Ensemble of Native α-Synuclein in Solution as Determined by Short-Distance Crosslinking Constraint-Guided Discrete Molecular Dynamics Simulations. PLoS Comput. Biol. 2019, 15 (3). https://doi.org/10.1371/journal.pcbi.1006859.

(47) Han, J. Y.; Choi, T. S.; Kim, H. I. Molecular Role of Ca2+ and Hard Divalent Metal Cations on Accelerated Fibrillation and Interfibrillar Aggregation of α-Synuclein. Sci. Rep. 2018, 8 (1), 1895. https://doi.org/10.1038/s41598-018-20320-5.

(48) Uversky, V. N.; Li, J.; Fink, A. L. Evidence for a Partially Folded Intermediate in α-Synuclein Fibril Formation. J. Biol. Chem. 2001, 276 (14), 10737–10744. https://doi.org/10.1074/jbc.M010907200.

(49) Bernstein, S. L.; Liu, D.; Wyttenbach, T.; Bowers, M. T.; Lee, J. C.; Gray, H. B.; Winkler, J. R. α-Synuclein: Stable Compact and Extended Monomeric Structures and PH Dependence of Dimer Formation. J. Am. Soc. Mass Spectrom. 2004, 15 (10), 1435–1443. https://doi.org/10.1016/J.JASMS.2004.08.003.

(50) Frimpong, A. K.; Abzalimov, R. R.; Uversky, V. N.; Kaltashov, I. A. Characterizationof Intrinsically Disordered Proteins with Electrospray Inization Mass Spectrometry: Conformationl Heterogeneity of α-Synuciein. Proteins Struct. Funct. Bioinforma. 2010, 78 (3), 714–722. https://doi.org/10.1002/prot.22604.

(51) Wu, K.-P.; Weinstock, D. S.; Narayanan, C.; Levy, R. M.; Baum, J. Structural Reorganization of α-Synuclein at Low PH Observed by NMR and REMD Simulations. J. Mol. Biol. 2009, 391 (4), 784–796. https://doi.org/10.1016/j.jmb.2009.06.063.

(52) Levitan, K.; Chereau, D.; Cohen, S. I. A. A.; Knowles, T. P. J. J.; Dobson, C. M.; Fink, A. L.; Anderson, J. P.; Goldstein, J. M.; Millhauser, G. L. Conserved C-Terminal Charge Exerts a Profound Influence on the Aggregation Rate of α-Synuclein. J. Mol. Biol. 2011, 411 (2), 329– 333.

(53) Illes-Toth, E.; Rempel, D. L.; Gross, M. L. Pulsed Hydrogen-Deuterium Exchange Illuminates the Aggregation Kinetics of α-Synuclein, the Causative Agent for Parkinson’s Disease. ACS Chem. Neurosci. 2018, 9 (6), 1469–1476. https://doi.org/10.1021/acschemneuro.8b00052.

(54) Del Mar, C.; Greenbaum, E. A.; Mayne, L.; Englander, S. W.; Woods, V. L. Structure and Properties of Alpha-Synuclein and Other Amyloids Determined at the Amino Acid Level. Proc. Natl. Acad. Sci. U. S. A. 2005, 102 (43), 15477–15482. https://doi.org/10.1073/pnas.0507405102.

(55) Kumar, H.; Singh, J.; Kumari, P.; Udgaonkar, J. B. Modulation of the Extent of Structural Heterogeneity in α-Synuclein Fibrils by the Small Molecule Thioflavin T. J. Biol. Chem. 2017, 292 (41), 16891–16903. https://doi.org/10.1074/jbc.M117.795617.

(56) Bertoncini, C. W.; Jung, Y.-S.; Fernandez, C. O.; Hoyer, W.; Griesinger, C.; Jovin, T. M.; Zweckstetter, M. Release of Long-Range Tertiary Interactions Potentiates Aggregation of Natively Unstructured α-Synuclein. Proc. Natl. Acad. Sci. 2005, 102 (5), 1430–1435. https://doi.org/10.1073/pnas.0407146102.

(57) Murray, I. V. J.; Giasson, B. I.; Quinn, S. M.; Koppaka, V.; Axelsen, P. H.; Ischiropoulos, H.; Trojanowski, J. Q.; Lee, V. M. Y. Role of α-Synuclein Carboxy-Terminus on Fibril Formation in Vitro. Biochemistry 2003, 42 (28), 8530–8540. https://doi.org/10.1021/bi027363r.

(58) Meuvis, J.; Gerard, M.; Desender, L.; Baekelandt, V.; Engelborghs, Y. The Conformation and the Aggregation Kinetics of α-Synuclein Depend on the Proline Residues in Its C-Terminal Region. Biochemistry 2010, 49 (43), 9345–9352. https://doi.org/10.1021/bi1010927.

(59) El Turk, F.; De Genst, E.; Guilliams, T.; Fauvet, B.; Hejjaoui, M.; Di Trani, J.; Chiki, A.; Mittermaier, A.; Vendruscolo, M.; Lashuel, H. A.;, et al. Exploring the Role of Post-Translational Modifications in Regulating α-Synuclein Interactions by Studying the Effects of Phosphorylation on Nanobody Binding. Protein Sci. 2018, 27 (7), 1262–1274. https://doi.org/10.1002/pro.3412.

(60) Nielsen, M. S.; Vorum, H.; Lindersson, E.; Jensen, P. H. Ca2+ Binding to ??-Synuclein Regulates Ligand Binding and Oligomerization. J. Biol. Chem. 2001, 276 (25), 22680–22684. https://doi.org/10.1074/jbc.M101181200.

(61) Kessler, J. C.; Rochet, J. C.; Lansbury, P. T. The N-Terminal Repeat Domain of α-Synuclein Inhibits β-Sheet and Amyloid Fibril Formation. Biochemistry 2003, 42 (3), 672–678. https://doi.org/10.1021/bi020429y.

(62) Shvadchak, V. V; Subramaniam, V. A Four-Amino Acid Linker between Repeats in the α-Synuclein Sequence Is Important for Fibril Formation. Biochemistry 2014, 53 (2), 279–281. https://doi.org/10.1021/bi401427t.

(63) Shaykhalishahi, H.; Gauhar, A.; Wördehoff, M. M.; Grüning, C. S. R.; Klein, A. N.; Bannach, O.; Stoldt, M.; Willbold, D.; Härd, T.; Hoyer, W. Contact between the Β1 and Β2 Segments of α-Synuclein That Inhibits Amyloid Formation. Angew. Chemie Int. Ed. 2015, 54 (30), 8837–8840. https://doi.org/10.1002/anie.201503018.

(64) Mirecka, E. A.; Shaykhalishahi, H.; Gauhar, A.; Akgül, Ş.; Lecher, J.; Willbold, D.; Stoldt, M.; Hoyer, W. Sequestration of a β-Hairpin for Control of α-Synuclein Aggregation. Angew. Chemie Int. Ed. 2014, 53 (16), 4227–4230. https://doi.org/10.1002/anie.201309001.

(65) Agerschou, E. D.; Flagmeier, P.; Saridaki, T.; Galvagnion, C.; Komnig, D.; Heid, L.; Prasad, V.; Shaykhalishahi, H.; Willbold, D.; Dobson, C. M.;, et al. An Engineered Monomer Binding-Protein for α-Synuclein Efficiently Inhibits the Proliferation of Amyloid Fibrils. Elife 2019, 8. https://doi.org/10.7554/eLife.46112.

(66) Krasnoslobodtsev, A. V.; Volkov, I. L.; Asiago, J. M.; Hindupur, J.; Rochet, J. C.; Lyubchenko, Y. L. α-Synuclein Misfolding Assessed with Single Molecule AFM Force Spectroscopy: Effect of Pathogenic Mutations. Biochemistry 2013, 52 (42), 7377–7386. https://doi.org/10.1021/bi401037z.

(67) Brucale, M.; Sandal, M.; Di Maio, S.; Rampioni, A.; Tessari, I.; Tosatto, L.; Bisaglia, M.; Bubacco, L.; Samorì, B. Pathogenic Mutations Shift the Equilibria of α-Synuclein Single Molecules towards Structured Conformers. ChemBioChem 2009, 10 (1), 176–183. https://doi.org/10.1002/cbic.200800581.

(68) Wise-Scira, O.; Aloglu, A. K.; Dunn, A.; Sakallioglu, I. T.; Coskuner, O. Structures and Free Energy Landscapes of the Wild-Type and A30P Mutant-Type α-Synuclein Proteins with Dynamics. ACS Chem. Neurosci. 2013, 4 (3), 486–497. https://doi.org/10.1021/cn300198q.

(69) Bertoncini, C. W.; Fernandez, C. O.; Griesinger, C.; Jovin, T. M.; Zweckstetter, M. Familial Mutants of α-Synuclein with Increased Neurotoxicity Have a Destabilized Conformation. J. Biol. Chem. 2005, 280 (35), 30649–30652. https://doi.org/10.1074/jbc.C500288200.

(70) Bussell, R.; Eliezer, D. Residual Structure and Dynamics in Parkinson’s Disease-Associated Mutants of α-Synuclein. J. Biol. Chem. 2001, 276 (49), 45996–46003. https://doi.org/10.1074/jbc.M106777200.

(71) Eliezer, D.; Kutluay, E.; Bussell, R.; Browne, G. Conformational Properties of α-Synuclein in Its Free and Lipid-Associated States. J. Mol. Biol. 2001, 307 (4), 1061–1073. https://doi.org/10.1006/JMBI.2001.4538.

(72) Ferreon, A. C. M.; Gambin, Y.; Lemke, E. a; Deniz, A. a. Interplay of Alpha-Synuclein Binding and Conformational Switching Probed by Single-Molecule Fluorescence. Proc. Natl. Acad. Sci. U. S. A. 2009, 106 (14), 5645–5650. https://doi.org/10.1073/pnas.0809232106.

(73) Fusco, G.; Pape, T.; Stephens, A. D.; Mahou, P.; Costa, A. R.; Kaminski, C. F.; Kaminski Schierle, G. S.; Vendruscolo, M.; Veglia, G.; Dobson, C. M.;, et al. Structural Basis of Synaptic Vesicle Assembly Promoted by α-Synuclein. Nat. Commun. 2016, 7, 12563. https://doi.org/10.1038/ncomms12563.

(74) Fauvet, B.; Fares, M. B.; Samuel, F.; Dikiy, I.; Tandon, A.; Eliezer, D.; Lashuel, H. A. Characterization of Semisynthetic and Naturally N α-Acetylated α-Synuclein in Vitro and in Intact Cells: Implications for Aggregation and Cellular Properties of α-Synuclein. J. Biol. Chem. 2012, 287 (34), 28243–28262. https://doi.org/10.1074/jbc.M112.383711.

(75) Fauvet, B.; Lashuel, H. A. Semisynthesis and Enzymatic Preparation of Post-Translationally Modified α-Synuclein. In Methods in Molecular Biology; Humana Press, New York, NY, 2015; Vol. 1345, pp 3–20. https://doi.org/10.1007/978-1-4939-2978-8_1.

